# Additive genetic effects in interacting species jointly determine the outcome of caterpillar herbivory

**DOI:** 10.1101/2022.01.20.476992

**Authors:** Zachariah Gompert, Tara Saley, Casey Philbin, Su’ad A. Yoon, Eva Perry, Michelle E. Sneck, Joshua G. Harrison, C. Alex Buerkle, James A. Fordyce, Chris C. Nice, Craig Dodson, Sarah L. Lebeis, Lauren K. Lucas, Matthew L. Forister

## Abstract

Plant-insect interactions are common and important in basic and applied biology. Trait and genetic variation can affect the outcome and evolution of these interactions, but the relative contributions of plant and insect genetic variation and how these interact remain unclear and are rarely subject to assessment in the same experimental context. Here we address this knowledge gap using a recent host range expansion onto alfalfa by the Melissa blue butterfly. Common garden rearing experiments and genomic data show that caterpillar performance depends on plant and insect genetic variation, with insect genetics contributing to performance earlier in development and plant genetics later. Our models of performance based on caterpillar genetics retained predictive power when applied to a second common garden. Much of the plant genetic effect could be explained by heritable variation in plant phytochemicals, especially saponins, peptides, and phosphatidyl cholines, providing a mechanistic understanding of variation in the species interaction. We find evidence of polygenic, mostly additive effects within and between species, with consistent effects of plant genotype on growth and development across multiple butterfly species. Our results inform theories of plant-insect coevolution and the evolution of diet breadth in herbivorous insects and other host-specific parasites.

**Teaser summary:** The combined, additive effects of plant and insect genetic variation explain Melissa blue caterpillar growth and development on alfalfa plants.

## Introduction

A central challenge for the biological sciences is to understand the causes and consequences of trait variation within and among species. Experimental manipulations aimed at understanding the molecular basis of organismal variation have most often been done in settings stripped of all or most ecological context. This approach can be fruitful for simple traits, including some aspects of morphology (e.g., [1–4]), but is lacking when it comes to interspecific interactions that include the evolution of crop pests, emerging infectious diseases, and other host-parasite associations [5, 6].

Plants and herbivorous insects have contributed much to our understanding of the formation and persistence of interactions between hosts and parasites, in part because they are experimentally tractable but also because insects are the most diverse macroscopic organisms on the planet and their specialized feeding habits play a role in their diversification [7–10]. Yet classic studies of the molecular basis of plant-insect interactions have relied on candidate genes or targeted classes of phytochemical compounds (e.g., [11–13]). More recently, evolutionary geneticists have taken advantage of new technologies to explore the genetic basis of herbivory in a genomic context. With very few exceptions, these studies have focused on genetic variation in either herbivores or plants [14–17](but see [18]), but rarely both in the same study and never to our knowledge paired with modern metabolomic approaches that allow for untargeted discovery of influential compounds [19]. This leaves us with considerable uncertainty concerning the relative importance of heritable traits in herbivores and in plants for determining the outcome of plant-insect interactions. For example, particular genetic variants in an herbivore might be associated with increased feeding efficiency, but only when challenged with particular plant variants such as specific defensive metabolites or combinations of physical defenses [20]. However, without an understanding of the genetic architecture of both the herbivore physiology and the plant traits, the evolutionary trajectory of the system cannot be understood in the context of available theoretical models or forecast with respect to the evolution of defense in the plant or increased performance in the herbivore. We address this need using a recent host range expansion onto alfalfa by the Melissa blue butterfly, emphasizing the role of prediction when building an understanding of the functional genetic basis of a novel plant-insect interaction.

The Melissa blue butterfly (*Lycaeides melissa*) is widespread in western North America [21]. It exists in isolated populations associated with larval host plants in the legume family, including many species of *Astragalus* and *Lupinus* [22, 23]. The Melissa blue colonized alfalfa (*Medicago sativa*) after the plant was introduced to the western USA as a forage crop in the mid 1800s, and is now commonly found on naturalized (i.e., feral) alfalfa along roadsides and trails [22]. Melissa blue butterflies show evidence of adaptation to alfalfa, but this host plant remains inferior to known native hosts in terms of caterpillar development with cascading life history effects [24–26]. Alfalfa is phenotypically variable [27], and thus is not a homogeneous resource for Melissa blue butterflies. In particular, phenotypic variation among naturalized alfalfa populations, including phytochemical variation, affects Melissa blue caterpillar growth and host patch occupancy [23, 28, 29]. However, it is unclear how much of this phenotypic variation has a genetic basis. Moreover, as is true for other plant-insect interactions, the relative contributions of plant (alfalfa) and insect (Melissa blue) genetic variation to the outcome of the interaction is unexplored, including whether growth and successful development from caterpillar to adult is influenced by additive or epistatic genetic variation in the interacting species.

Here, we use multiple common garden rearing experiments combined with multilocus genetic mapping and genomic prediction to build and test models that quantify the relative effects and interactions of alfalfa and Melissa blue genetic variation on caterpillar performance (i.e., growth and survival). We specifically test the following alternative hypotheses: (i) caterpillar performance is primarily affected by Melissa blue genetic variation and architecture, (ii) caterpillar performance is primarily affected by genetic variation and architecture in the host plant, (iii) the genetics of the interacting species have similar effects on caterpillar performance and combine additively, (iv) the genetics of the interacting species have similar effects on caterpillar performance and combine epistatically, and (v) the null hypothesis that neither Melissa blue nor alfalfa genetic variation has an appreciable effect on caterpillar performance (Fig. 1). Genetic mapping of 1760 plant traits, including 1750 phytochemical metabolites, contributes to testing these hypotheses and also allows us to probe the functional basis of plant-genetic effects on caterpillar performance. Finally, we conduct complementary rearing experiments to test the consistency of plant genetic effects (i.e., their lack of interaction with herbivore genetics) across butterfly populations and species.

**Figure 1:**
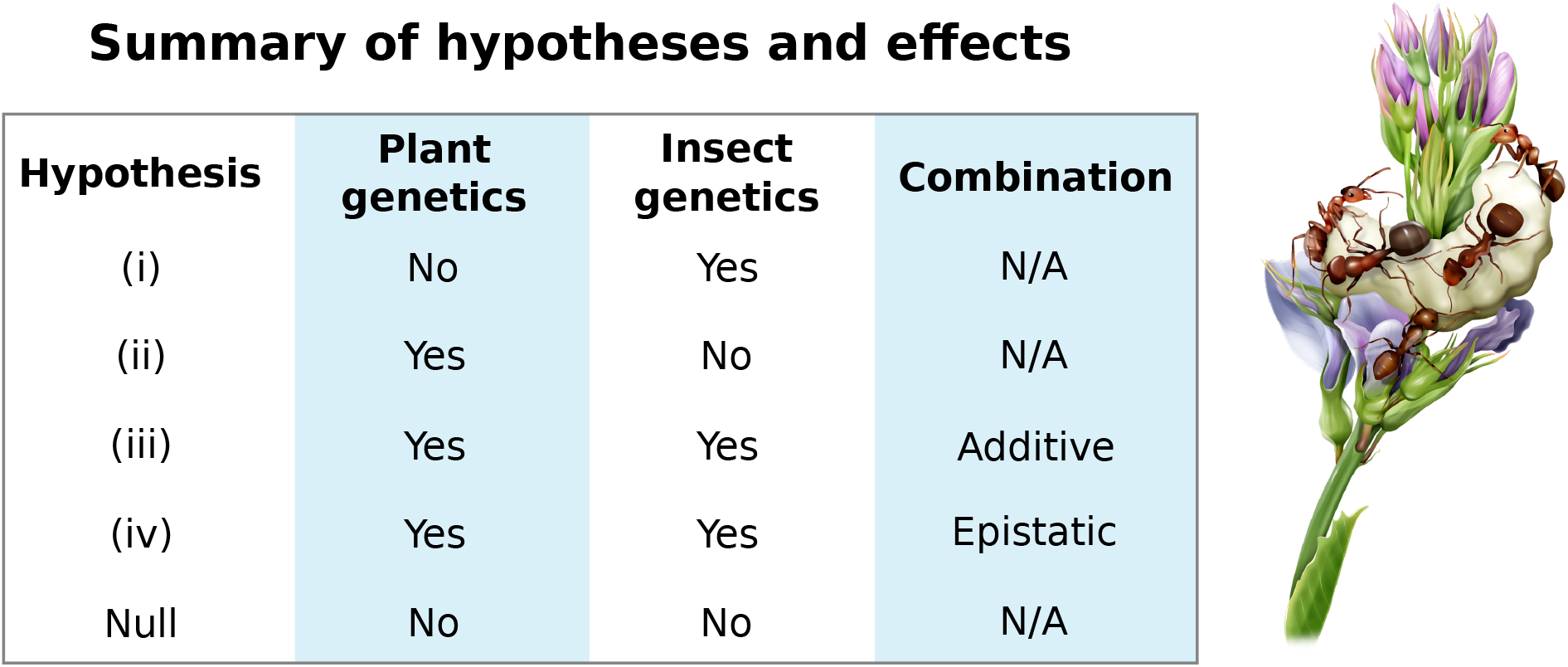
Main hypotheses tested about the contribution of plant and insect genetics to caterpillar performance: (i) caterpillar performance is primarily affected by insect (*L. melissa*) genetics, (ii) caterpillar performance is primarily affected by plant (*M. sativa*) genetics, (iii) the genetics of the interacting species have similar effects on caterpillar performance and combine additively, (iv) the genetics of the interacting species have similar effects on caterpillar performance and combine epistatically, and (v) the null hypothesis that neither insect or plant genetic variation have an appreciable effect on caterpillar performance. The illustration (by R. Ribas) shows a *L. melissa* caterpillar feeding on alfalfa, while being tended by ants; additional biotic or abiotic factors, such as the presence of mutualistic ants, also affect caterpillar performance in the wild [23] but are not a component of this study.

## Results

### Overview of the primary common garden rearing experiment

We planted a common garden comprising 1080 alfalfa (*M. sativa*) plants at the Greenville Experimental Farm near Logan, UT (41.765° N, 111.814° W) in 2018 (Fig. S1a). Seeds for this garden were collected from 11 naturalized (i.e., feral) *M. sativa* sites in the western USA, including five sites where *L. melissa* butterflies are found (Fig. 2a). Caterpillars for the experiment were sourced from six sites by obtaining eggs from gravid *L. melissa* females in 2019. We detected substantial genetic variation and only subtle genetic differentiation among the source locations for alfalfa (161,008 SNPs, mean expected heterozygosity = 0.168, F_ST_ = 0.029) and for *L. melissa* (63,194 SNPs, mean expected heterozygosity = 0.065, F_ST_ = 0.045) (Figs. 2b, S2). The main rearing experiment was conducted in summer 2019. For this experiment, caterpillars were reared individually on each of the 1080 alfalfa plants. Rearing was done in a growth chamber, with caterpillars fed fresh leaf tissue as needed. In this experiment, 26.1% of the caterpillars survived to pupation and 14.1% survived to eclose as adults (mean survival time = 21.8 days) (Fig. 3a). Mean *L. melissa* weights were 2.94 mg (SD = 2.13) at 8 days, 12.7 mg (SD = 7.71) at 14 days, and 20.0 mg (SD = 7.21) at pupation. Weight and survival metrics of performance were positively correlated, including, 8-day weight vs. 14-day survival (Pearson *r* = 0.0916, 95% confidence interval [CI] = 0.0237–0.159), 14-day weight vs. survival to pupation (*r* = 0.472, 95% CI = 0.416–0.525), and pupal weight vs. survival to eclosion (*r* = 0.449, 95% CI = 0.342–0.545) (Fig. 3b). Past work has shown that weight and lifetime fecundity are highly correlated in *L. melissa* [24].

**Figure 2:**
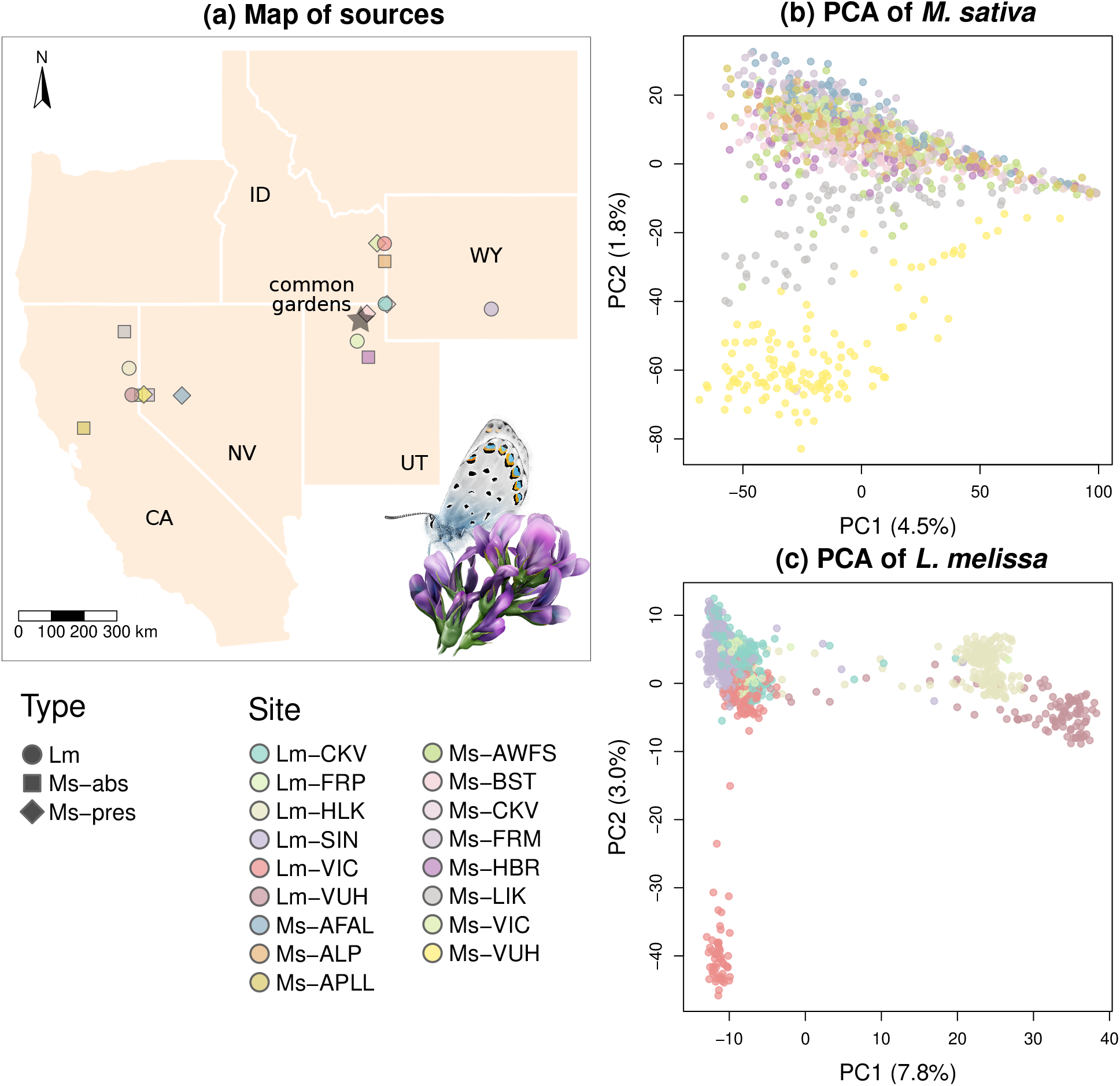
(a) Map of plant (*M. sativa*) and insect (*L. melissa*) common garden source populations. Symbol shapes denote source type–Lm = *L. melissa*, Ms-abs = *M. sativa* site without *L. melissa* butterflies, and Ms-pres = *M. sativa* site with *L. melissa* butterflies, and are colored to indicate different populations within taxa. The inset illustration shows an adult *L. melissa* perched on *M. sativa* (illustration by R. Ribas). (b) Ordination of genetic variation via principal component analysis (PCA) for the *M. sativa* common garden plants. (c) Ordination of genetic variation via PCA for the *L. melissa* caterpillars from the rearing experiment. Points in (b) and (c) denote individual plants or caterpillars and are colored to match the map (a).

**Figure 3:**
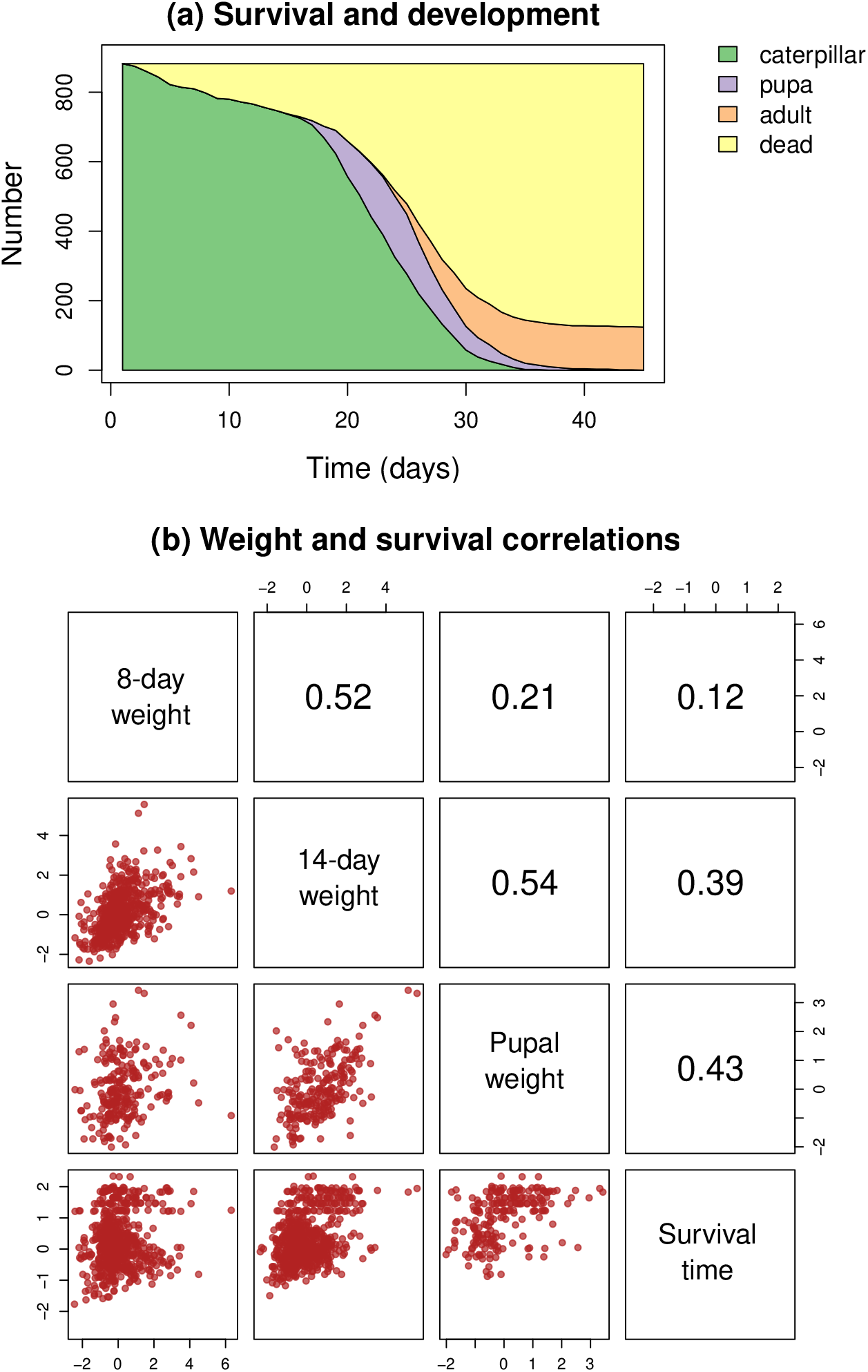
(a) Plot shows survival and development of *L. melissa* over the course of the rearing experiment. Colored regions denote the number of individuals that were living caterpillar, pupa, adults or dead at each day post hatching. (b) Plots show pairwise correlations between *L. melissa* performance traits. Scatterplots are shown in the lower-triangle panels–each point denotes one individual–and Pearson correlations are reported in the corresponding upper triangle panels. Traits are given along the diagonal panels: 8-day weight, 14-day weight, pupal weight, and truncated survival time. Scatterplots and Pearson correlations are based on residuals after controlling for confounding environmental effects (see Methods for details).

### Plant and caterpillar genetic variation affect performance

Using multilocus genome-wide association methods, we found evidence that both *M. sativa* and *L. melissa* genetic variation contributed to caterpillar performance in the common garden rearing experiment (Fig. 4a), consistent with our hypotheses (iii) and (iv) (Fig. 1). Specifically, *M. sativa* genetics (161,008 SNPs) explained between 2% (survival to 8 days) and 36% (14-day weight) of the variation in performance (mean across traits = 17%), and *L. melissa* genetics (63,194 SNPs) explained 5% (weight at pupation and survival to pupation) to 29% (8-day weight) of the variation in the same nine caterpillar performance measures (mean = 15%) (values denote point estimates of the percent variance explained, PVE; see Table S2 for credible intervals; cross-validation results are shown in the next section). Caterpillar genetics contributed more to performance metrics from early development (e.g., 8-day weight and survival to 8 and 14 days), whereas plant genetics mattered more for later development (e.g., 14-day weight, pupal weight, and survival to pupation and adult), resulting in a trend towards a negative relationship between caterpillar and plant genetic contributions across traits (Pearson *r* = −0.52, 95% CI = −0.88 to 0.22, *P* = 0.15). We detected mostly positive genetic correlations among performance traits (Fig. 4b), with similar but not identical genetic correlations calculated from *M. sativa* and *L. melissa* polygenic scores (Pearson correlation between *M. sativa* and *L. melissa* genetic correlations, *r* = 0.80, 95% CI = 0.63 to 0.89, *P* = 5.923e^-9^).

**Figure 4:**
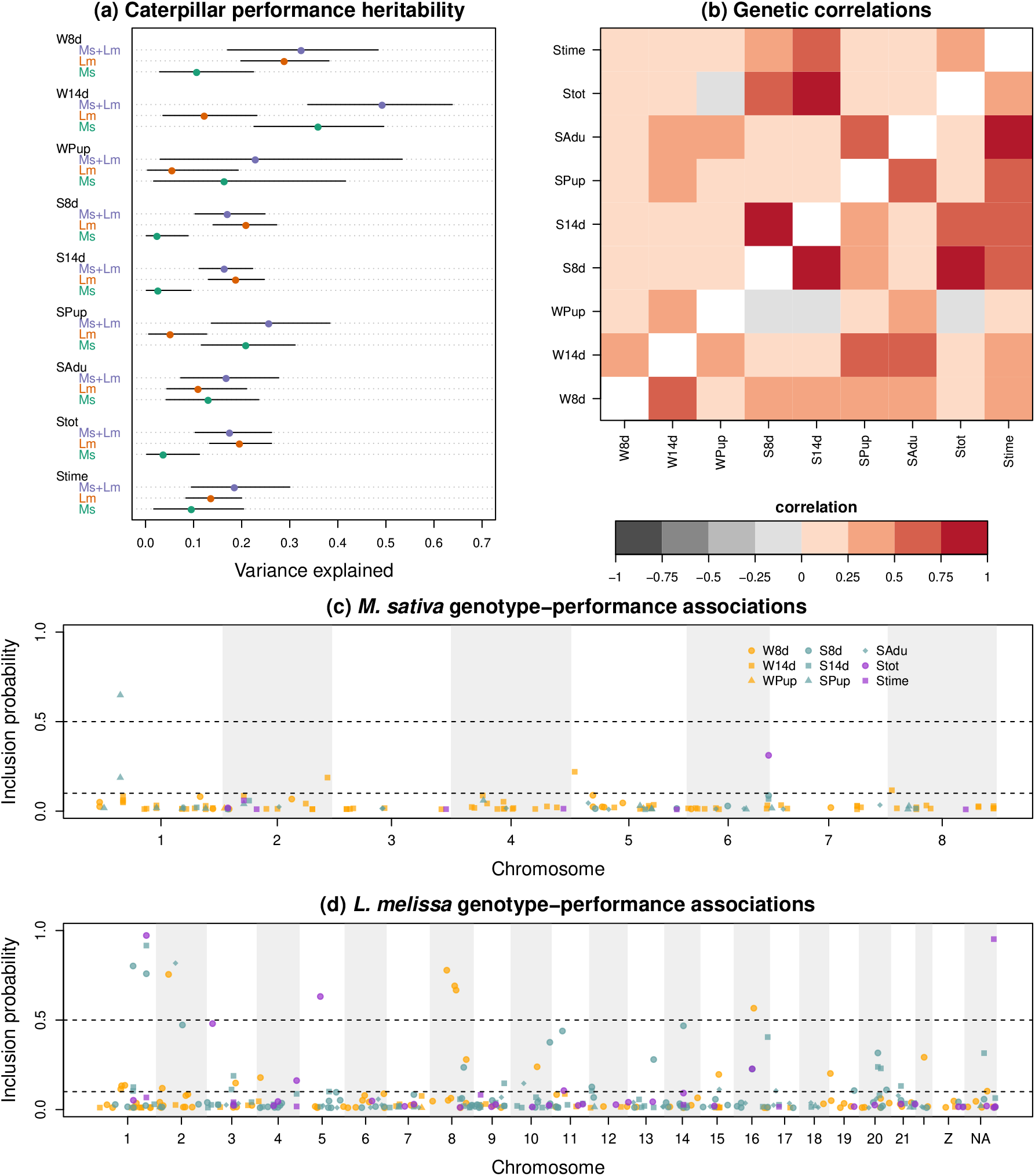
Genetic mapping of caterpillar performance. (a) Dotchart shows Bayesian estimates of the proportion of trait variation explained by *M. sativa* genetics (Ms), *L. melissa* genetics (Lm), or both combined (Ms+Lm) for each caterpillar performance trait; W8d = 8-day weight, W14d = 14-day weight, Wpup = pupal weight, S8d = 8-day survival, S14d = 14-day survival, SPup = survival to pupation, SAdu = survival to adult, Stot = total survival time, and Stime = (truncated) survival time. Points and horizontal lines denote point estimates (posterior medians) and 95% equal-tail probability intervals, respectively. (b) Heatmap shows genetic correlations between pairs of caterpillar performance traits based on *M. sativa* genetics (lower triangle) or *L. melissa* genetics (upper triangle). Manhattan plots in (c) and (d) shown posterior inclusion probabilities (PIPs) for genotype-performance associations based on *M. sativa* and *L. melissa* SNPs, respectively. Points denote SNPs with different colors and symbols for different performance traits. Only SNPs with PIPs ≥ 0.01 are depicted. Horizontal lines at PIPs of 0.1 and 0.5 are included for reference.

Mapping results suggested mostly a polygenic basis for the performance traits, with point estimates of > 10 loci affecting most traits (Tables S2, S3, S4; Fig. 4c,d), but with more evidence of specific SNPs strongly associated with performance in *L. melissa*. This included ten SNPs with posterior probabilities of association (i.e., posterior inclusion probabilities) > 0.5 with at least one performance trait (Fig. 4d, Table S5). Some of these SNPs were in or near (<20 kbps) genes with biologically plausible functions for affecting performance, such as *MSP-300*, *Lipase member H* and *Juvenile hormone acid O-methyltransferase*, all of which were associated with 8-day weight. For example, *MSP-300* affects muscle development and muscle-ectoderm attachment in *Drosophila* [30]. Insect lipases metabolize fats, are expressed in gut tissue, and can affect survival and reproductive capacity in insects; *Lipase member H* in particular has further been associated with viral resistance in the moth *Bombyx mori* [31,32]. *Juvenile hormone acid O-methyltransferase* is involved in juvenile hormone biosynthesis and thus in the regulation of insect growth and development, especially metamorphosis [33, 34]. A single *M. sativa* SNP was strongly associated with survival to pupation (posterior inclusion probability > 0.5; chromosome 1, position = 12,930,966 bp). This SNP was found in a gene encoding *TOM1-like protein 9* and was within 30 kbps of six additional genes, including two genes with known links to plant-insect interactions: *dentin sialophosphoprotein*, which is associated with soybean compensatory growth after cutworm herbivory [35], and *photosystem I reaction center subunit psaK*, which has been mechanistically linked to tolerance to aphids and aphid feeding preference in *Arabidopsis* [36] (Table S6).

We repeated the genetic mapping approach using a combined data set of both *M. sativa* and *L. melissa* genetic loci (i.e., the combined 224,202 SNPs). The combined data set generally explained more of the variation in caterpillar performance, 17% to 49% (mean = 24%), than either *M. sativa* or *L. melissa* genetic loci alone. Moreover, the combined variation explained for each performance trait was well described by a model where the variance explained separately by plant and caterpillar genetics combined additively, consistent with our hypothesis (iii) (Fig. 1). Specifically, in a linear regression model, the percent variance in performance traits explained by plant and caterpillar genetics separately explained 97% of the variation in the estimates of the variance explained by the combined genetic data sets (linear regression, *β_plant_* = 1.17, *P* = 6.6*e*^-6^, *β_caterpillar_* = 0.80, *P* = 0.00037, *r*^2^ = 0.97) (Fig. S4).

### Predicting caterpillar performance from plant and caterpillar genotype

We next showed that our genotype-phenotype models were moderately successful at predicting caterpillar performance. This is relevant both for validating these models and for demonstrating their potential utility and limitations in making predictions about effects and evolutionary trajectories in nature. Specifically, genomic predictions of performance from 10-fold cross validation exhibited statistically significant positive correlations with observed performance values for three out of ten performance traits for *M. sativa* genetics, five out of ten traits for *L. melissa* genetics, and six out of ten traits for *M. sativa* and *L. melissa* genetics combined (Fig. 5a). Especially pronounced positive correlations between observed and predicted performance were detected for 14-day weight based on *M. sativa* genetics and 8-day and 14-day survival based on *L. melissa* genetics. More generally, our ability to predict performance traits was well explained by our estimates of heritability (i.e., PVE): we calculated Pearson correlations of 0.89 (95% CI = 0.55 to 0.98, *P* = 0.0013) and 0.62 (95% CI = −0.073 to 0.91, *P* = 0.074) between PVE estimates and the correlation between observed and predicted traits for *M. sativa* and *L. melissa* genetics, respectively (Fig. 5b). In other words, we better predicted caterpillar performance for the performance traits that were more heritable.

**Figure 5:**
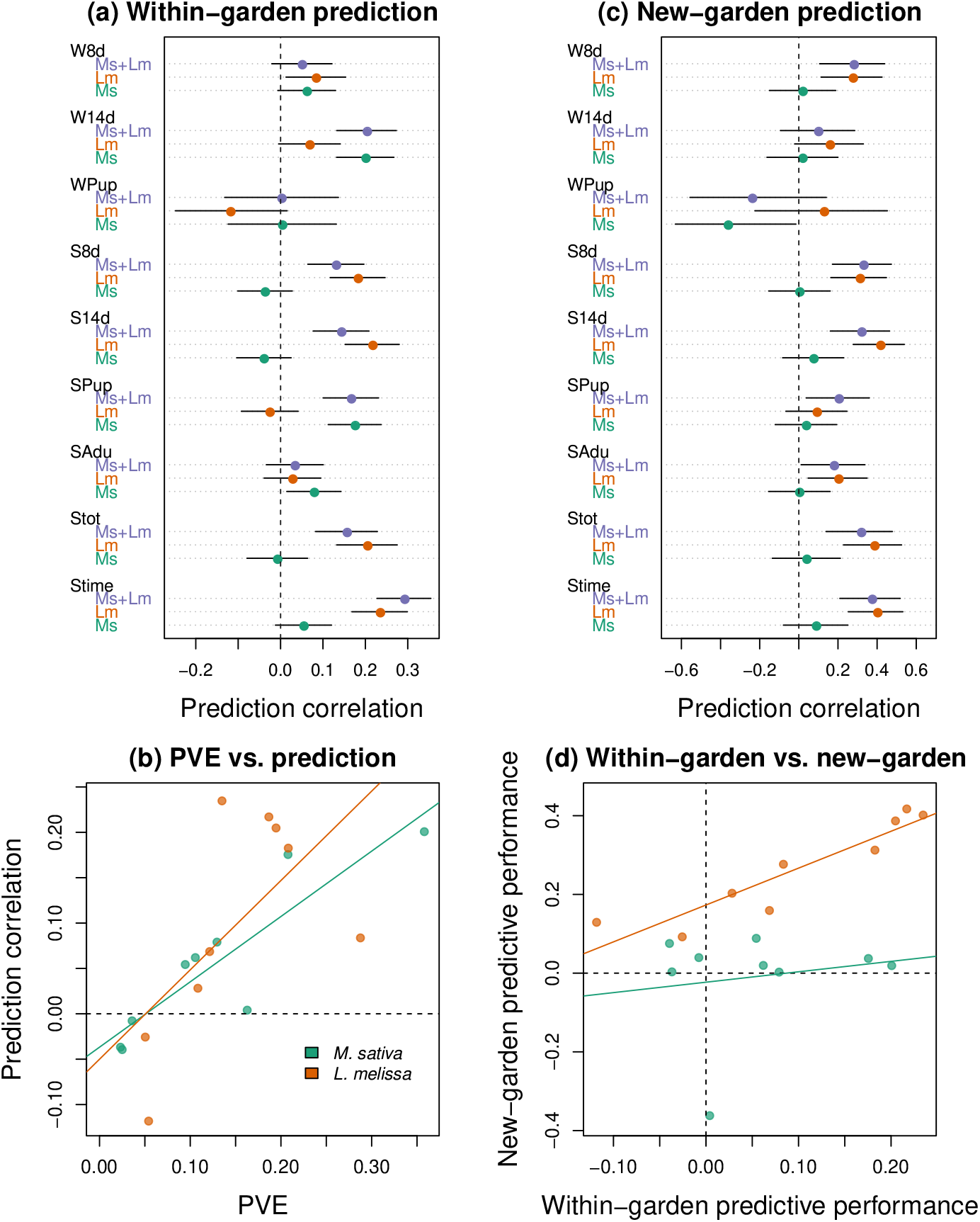
Genomic prediction of caterpillar performance. (a) Dotchart shows Pearson correlations between cross-validation genomic predictions of phenotypes and the observed values based on *M. sativa* genetics (Ms), *L. melissa* genetics (Lm), or both combined (Ms+Lm) for each caterpillar performance trait; W8d = 8-day weight, W14d = 14-day weight, Wpup = pupal weight, S8d = 8-day survival, S14d = 14-day survival, SPup = survival to pupation, SAdu = survival to adult, Stot = total survival time, and Stime = (truncated) survival time. Points and horizontal lines denote point estimates (posterior medians) and 95% equal-tail probability intervals, respectively. For example, a large value on the x axis indicates a high correlation between observed performance values and predictions from genotype based on cross validation. (b) Scatterplot of the proportion of variation explained by genetics (PVE) versus the Pearson correlation of genomic predictions from (a). Each point denotes a trait and is colored to indicate values from *M. sativa* or *L. melissa* genetics. Colored lines are best fits from ordinary linear regression, and a dashed line denotes the 0 value on the y-axis. (c) Dotchart similar to (a), but for genomic predictions of phenotypes in a second common garden (the Gene Miller Life Science Garden) based on the models fit from the main garden. (d) Scatterplot of correlations between observed caterpillar performance trait values and genomic predictions of these values using cross-validation within the main garden versus prediction for samples in the Gene Miller Life Science Garden based on the models fit for the main garden. Symbols, colors and lines are as defined for (b).

Having demonstrated moderate predictive power within the main common garden, we next asked whether genotype-phenotype models estimated from this garden could successfully predict *L. melissa* performance for additional caterpillars fed *M. sativa* from a second, smaller common garden (the Gene Miller Life Science Garden; *N* = 180 plants) (Fig. S1). This second garden, planted in 2018 on the Utah State University campus ~2.5 km from the Greenville Experimental Farm garden, included plants from six of the 11 *M. sativa* source sites and caterpillars from each of the sites used in the main experiment. Survival rates for caterpillars reared on plants from this garden were similar to those reared on plants from the main garden (Fig. S3). Predictive performance for the second garden differed notably for *M. sativa* versus *L. melissa* genotype-phenotype models, with statistically significant positive correlations between observed and predicted trait values in the new garden for only one trait for *M. sativa* genetics versus six of the ten performance traits for *L. melissa* genetics (Fig. 5c). Predictions for the combined data set were similar to those based on *L. melissa* genetics alone. Consistent with these patterns, estimates of PVE from the main garden explained predictive power for *L. melissa* genetics (Pearson *r* = 0.93, 95% CI = 0.68 to 0.98), but not *M. sativa* genetics (*r* = 0.17, 95% CI = −0.56 to 0.75). Thus, unmeasured environmental differences likely limit our ability to predict performance from plant genetics across gardens to a much greater extent than for caterpillar genetics (plant growth environments differed, but caterpillar rearing environments did not) despite these gardens being separated by only ~2.5 km. Differences in the exact genetic composition of the two gardens could add to this effect.

### Genetic associations with plant traits explain the plant genetic contribution to caterpillar performance

Having shown that plant genetic variation affects caterpillar performance, we now focus on the Greenville Experimental Farm (see Fig. S1a) to identify components of the functional basis of the documented plant-genetic effects. This also allowed us to further test for additive versus epistatic interactions between plant and caterpillar genotypes on caterpillar performance (see our hypotheses (iii) versus (iv) in Fig. 1). We first determined the extent to which genetic loci associated with caterpillar performance were also associated with other plant traits, including potential plant vigor or defense traits [16]. Such an association would be consistent with the hypothesis that these traits, or other genetically correlated traits, constitute the mechanisms by which plant genotype affects caterpillar performance. To do this, we measured and mapped 1760 plant traits in the Greenville Experimental Farm garden using the same multilocus mapping approach and *M. sativa* SNP data set described above. The traits included plant height, leaf length, leaf width, leaf area, leaf shape, leaf weight, specific leaf area, leaf toughness, trichome density, levels of herbivory on the plants in the field, and 1750 plant chemistry metabolites, which were quantified and characterized using liquid-chromatography combined with mass spectrometry (LC-MS; similar to [23, 29]).

We documented genetic variation affecting most of the plant traits, with mean PVEs of 20.5% for the non-chemical traits (minimum = 5.6%, maximum = 38.7%) and 10.9% (310 traits > 20% and 20 > 50%) for the 1750 chemical traits (Table S7). Additionally, in the main Greenville Experimental Farm common garden, the distribution of PVE for the 1750 chemical traits differed markedly from that for 1750 matched, randomized traits, consistent with a clear genetic contribution to this variation in leaf metabolites (Fig. S5).

Multiple plant traits, including chemical and non-chemical traits, exhibited genetic correlations with each caterpillar performance trait; in other words, plant trait polygenic scores were correlated with caterpillar performance polygenic scores when inferred from plant genetics (Figs. 6a,b, S6). However, because of the large number of measured traits and genetic correlations among the plant traits (Fig. S7), many of the genetic correlations between plant traits and caterpillar performance were likely redundant. Thus, to identify the combined subset of traits most strongly predictive of caterpillar performance, we next fit a LASSO penalized regression model for the polygenic scores of each caterpillar performance trait (based on plant genetics) as a function of the polygenic scores for the 1760 plant traits. These models explained 41 to 80% of the variation in the caterpillar performance scores (mean = 69.2%, cross-validation predictive *r*^2^ ranged from 0.39 to 0.76) (Table S8, Fig. 6c). On average 260 of the 1760 traits were retained in these models (i.e., given non-zero regression coefficients), with a range of 117 (survival time) to 347 (8-day survival) traits (Figs. 6d,e and S8). Both chemical and non-chemical traits were retained in the models. Non-chemical traits with the biggest effects included a positive effect of plant height on 14-day weight (*β* = 0.037), positive effects of trichome density (*β* = 0.036) and specific leaf area (*β* = 0.031) on survival to adulthood, and a negative effect of leaf toughness on survival to adulthood (*β* = −0.34). Consistent with a previous phenotypic assay of caterpillar performance and plant metabolomic variation in this system [29], top chemical traits included several saponins, including saponins associated with reduced caterpillar weight (Quillaic acid) and increased survival (e.g., Medicagenic acid 3-O-beta-D-glucoside) (Tables S10, S11). The flavonoid glycoside Tricin 7-glucoside was associated with reduced survival, whereas several peptides (e.g., MESA.583 = C_13_H_18_O, MESA.615 = C_23_H_43_N_7_O_7_, MESA.849 = C_14_H_19_NO_3_) were associated with reduced weight or survival (Tables S10, S11).

**Figure 6:**
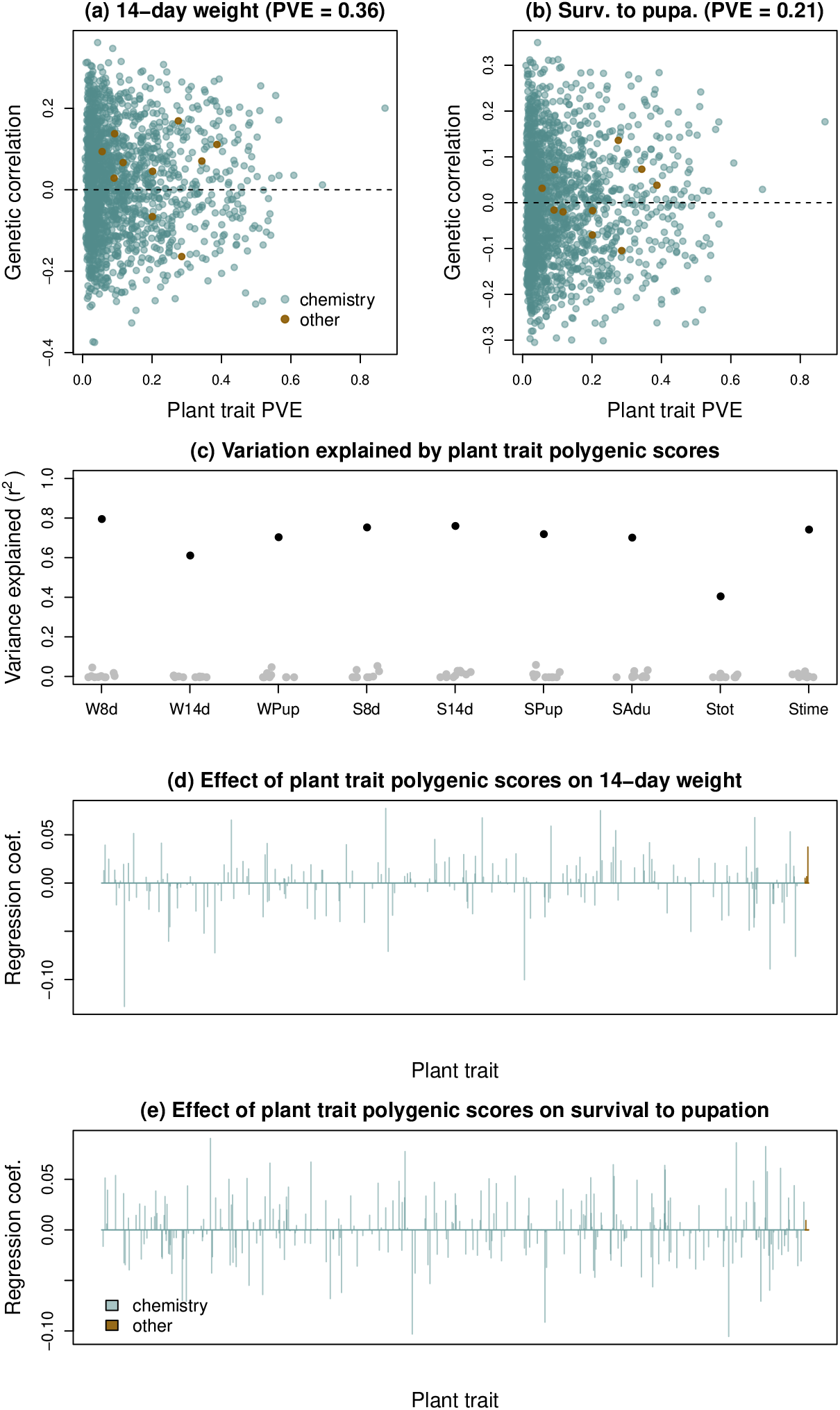
Associations between plant trait polygenic scores and caterpillar performance polygenic scores. Scatterplots show genetic correlations between plant chemistry and other plant traits and 14-day caterpillar weight (a) and survival to pupation (b) inferred from plant genetics as a function of the proportion of plant trait variation explained by genetics (PVE). A dashed horizontal line denotes a genetic correlation of zero. Panel (c) shows the variance explained by lasso regression models of caterpillar performance polygenic scores estimated from plant genetics as a function of polygenic scores for 1750 plant chemistry traits and 10 non-chemistry traits. Black dots denote inferred values of *r*^2^ and gray dots show similar estimates using randomized plant trait polygenic scores (10 random data sets each). Panels (d) and (e) show standardized regression coefficients from the lasso models for 14-day weight (d) and survival to pupation (e).

Compared to predicting polygenic scores for caterpillar performance, our ability to predict caterpillar performance at the phenotypic level from plant-trait polygenic scores was notably reduced (Table S8, Figs. S9, S10). This was expected as plant genetics only explained a modest proportion of the variation in performance and thus the ability to explain variation in these traits (not just polygenic scores) was necessarily capped by performance-trait heritabilities. Still, when considering all performance traits together, plant trait polygenic scores explained more of the trait variation than expected by chance (Fisher combined test, *χ*^2^ = 34.42, df = 18, *P* = 0.011). This signal was driven primarily by association of plant traits with 8 and 14 day weight and survival to pupation and eclosion.

Lastly, we determined the extent to which the association of plant trait polygenic scores with caterpillar performance polygenic scores (both inferred from plant genetics) was affected by *L. melissa* genotype. Such an interaction would suggest caterpillar performance is affected by epistatic interactions between *M. sativa* and *L. melissa* genotypes, as predicted by our hypothesis (iv) (Fig. 1). We used principal component (PC) scores from the first four principal components of the *L. melissa* genotype matrix as summaries of *L. melissa* genotype. We then fit LASSO penalized regression models for caterpillar performance polygenic scores as a function of these PC scores, plant trait polygenic scores, and interactions between each plant trait polygenic score and each of the four PCs. This allowed us to test for epistasis at the level of plant morphology and phytochemistry polygenic scores from *M. sativa* and four axes of *L. melissa* genetic background and thereby avoid the lack of power that would be associated with exhaustively testing SNP-SNP interactions. We found no evidence of epistasis between *M. sativa* and *L. melissa* affecting caterpillar performance. Specifically, including these interaction terms in the models actually reduced the variance explained by the LASSO models (Table S9) and the interaction terms were retained less frequently in the models than the non-interaction terms (Figs. S9, S11). We obtained similar results when fitting models for caterpillar performance trait values rather than polygenic scores, with a smaller proportion of interaction terms retained in the model for most traits (Figs. S13 and S14) and no overall increase in variance explained by models with versus without interactions (i.e., the variance explained in 14-day weight doubled, but the variance explained in 8-day weight was halved, and there was no detectable general increase in variance explained across traits) (Fig. S15). Thus, these results support our hypothesis (iii) with additive contributions of plant and caterpillar genetics (Fig. 1).

### Plant and caterpillar genetics have consistent effects on performance

We conducted two additional experiments to determine the extent to which genetic differences among *M. sativa* plants or populations had consistent affects on caterpillar performance for different butterfly populations and species. This constitutes another test of additivity versus epistasis for plant and insect genotypes (our hypotheses (iii) versus (iv) in Fig. 1) and of the potential for our findings to provide general predictions beyond our main study populations. In the first of these experiments, *L. melissa* caterpillars from four populations were reared on greenhouse-grown *M. sativa* sourced from six sites (Table S12). Two additional butterfly species, *Colias eurytheme* (a legume specialist) and *Vanessa cardui* (a generalist that rarely feeds on alfalfa), were reared on these same plants. Caterpillars were fed leaf tissue from multiple individual plants, but each caterpillar was given plants from a single source population. Survival rates were highest for *C. eurytheme*, followed by *L. melissa* and lastly *V. cardui* (Fig. S16). Plant population (here used as a proxy for plant genotype) explained ~3-10% of the variation in 8-day weight for each butterfly species, and 9-14% of the variation in 14-day weight, with larger effects in the butterfly species less-well adapted to *M. sativa* (Table S13). Caterpillar population explained a small but non-zero proportion of the variation in 8-day weight in *L. melissa* (this could not be assessed in the other species), but not a significant amount of variation in 14-day weight. Thus, consistent with our main results above, genetic differences among plant and caterpillar populations (caterpillar populations for *L. melissa* only) explained variation in caterpillar performance, with plant genetics mattering more for 14-day weight than 8-day weight and caterpillar genetics mattering more for 8-day weight than 14-day weight. Plant population and plant maternal family also explained variation in plant growth and development traits, consistent with our common garden results above (Table S7). Importantly, the effect of each plant population on caterpillar performance was remarkably consistent across *L. melissa* populations and even across different species, with moderate to large positive correlations (though not always significantly so) in the effect of each plant population on 8 and 14-day weight across all pairs of population and species (Fig. S17).

The final complementary experiment used the same three butterfly species: *L. melissa*, *C. eurytheme*, and *V. cardui*, but instead involved feeding each caterpillar leaf tissue from a single *M. sativa* plant from a third common garden near the University of Nevada (UNR Main Station in Reno, NV; Fig. S1). We used these data to ask whether the effect of plant genotype (here, individual plant) on caterpillar weight was consistent across species. We detected modest, positive pairwise correlations between the three species of caterpillars, suggesting a degree of similarity of plant genotypes that affect performance of these different herbivorous species (Fig. S18). Specifically, the correlations were as follows: *V. cardui* vs. *L. melissa r* = 0.33 (*P* = 0.015, t = 2.52, df = 52); *C. eurytheme* vs. *L. melissa r* = 0.43 (*P* = 0.0010, t = 3.48, df = 52); *C. eurytheme* vs. *V. cardui r* = 0.15 (P = 0.28, t = 1.08, df = 52). Thus, these two experiments combined with our main results show that genetic variation within *M. sativa* affects caterpillar performance across populations and species of butterflies in a remarkably consistent manner, consistent with the additivity hypothesis (hypothesis (iii) in Fig. 1).

## Discussion

From an ecological perspective, the greatest diversity of life is not counted in the number of species or other taxonomic units, but in the diversity of inter-specific interactions [37]. The ubiquity of plant-feeding insects has made them a focal point for understanding the evolution, persistence, and variability of interactions [38, 39]. In terms of variability, outcomes (e.g., caterpillar survival) could depend on genetic variation within each species and these genetic effects could compound additively or non-additively. Our results support the hypothesis that both plant (alfalfa) and insect (Melissa blue butterfly) genotype matter for caterpillar growth and survival, and that these contributions are mostly additive (our hypothesis (iii) in Fig. 1). These results are qualitatively similar to those reported in another study [18], which identified individual plant (*Arabidopsis thaliana*) and caterpillar (*Pieris rapae*) genes affecting caterpillar performance. The advance over previous work that we offer here is in quantitative, genomic prediction of caterpillar performance, which, in contrast to the identification of specific genes, provides a formal connection from trait genetics to models of evolution for quantitative traits [40]. We specifically demonstrated that the combined effects of plant and insect genotype explain a substantial proportion of variation in caterpillar growth and survival (17-49%), and that these mostly-additive effects can predict performance from genotypes in cross-validation analyses. Moreover, we were able to identify specific traits and phytochemicals associated with the plant contribution to performance, most notably plant size, and several saponins, peptides, and phosphatidyl cholines. Whereas some of these classes of chemicals (e.g., saponins) are best known as insect toxins or feeding-deterrents (e.g., [41–43]), our results suggest these classes include molecules with positive and negative effects on performance, consistent with other recent metabolomic work [23, 29]. We also found evidence that plant genotype had consistent effects on performance in multiple butterfly populations and species, including a second legume specialist (*C. eurytheme*) and a generalist ( *V. cardui*). This too is consistent with results from the only other similar study [18], which documented conserved changes in gene expression in response to herbivores across multiple plant and butterfly species. This consistency is relevant to the predictability and nature of the evolution of plant-insect interactions, as we discuss more below.

Our results have clear implications for the study of coevolution, which takes many forms and pertains to the formation of new species and new interactions [38]. Quantitative theories of coevolution have historically been dominated by gene-for-gene models, in which the fitness of a particular genetic variant in (for example) a parasite is conditioned on the presence of a specific gene in the host [20]. Evidence in support of gene-for-gene models has come mostly from plant-pathogen systems [20] (but see [44]). In contrast, diffuse models of coevolution relax some of the expectations for gene-by-gene interactions, and have been favored by researchers working with more macroscopic parasites, including herbivorous insects [45]. However, relevant investigations in plants and insects have mostly relied on experiments that contrast categories of individuals (strains or biotypes) rather than more comprehensive or continuous variation in genetically variable populations (reviewed in [11]), which has left the field with uncertainty regarding the most relevant theoretical context for the diversity of evolving plant-insect interactions. The results that we report are not consistent with the gene-for-gene model of coevolution, as the performance of our focal herbivore was both highly polygenic and successfully predicted without interactions between caterpillar and plant genotypes. Instead, our results suggest that genetic differences in plant quality and defense have similar effects regardless of insect genotype or even species.

Our results also shed light on the evolution of diet breadth and host use in herbivorous insects. Specifically, the finding of substantial heritable variation in the Melissa blue butterfly for growth and survival suggests that ongoing adaptation to alfalfa, which at present is a marginal host [24], is not constrained by a lack of genetic variation. This is consistent with earlier work on this system [26]. Likewise, alfalfa appears to harbor genetic variation to evolve traits that reduce the success of the Melissa blue even further, and this inference likely extends to other herbivores given the consistent effects of plant variation on other butterfly species reported here and on other herbivores in an observational study [23]. While the persistence of plant genetic variation affecting herbivores might be attributable to the age of these interactions (since most herbivores of alfalfa in North America are recent colonists), we suspect other factors are more important. First, the asymmetry in our predictions, with consistent caterpillar-genetic effects but not plant-genetic effects on performance between common gardens, suggests a major role for plasticity in the effect of plant genotype on caterpillar performance. This is not surprising given considerable evidence that biotic and abiotic environmental factors affect plant quality and plant defenses in alfalfa [28] and other plants [46], but does mean genetic variation in performance measured in the lab and common garden might not strongly predict effects in specific natural populations [47]. Moreover, other biotic and abiotic factors could contribute more to caterpillar growth and survival in the wild, and some of these could interact with plant genotype. For example, recent work has shown that the abundance of ants, which tend Melissa blue caterpillars and thereby reduce the threat from enemies (see image in Fig. 1), greatly increases caterpillar survival and population persistence on alfalfa, with ant abundance indirectly affected by alfalfa phytochemistry [23, 48]. In contrast to the complexity of plant effects, the more consistent effects of caterpillar genetic variants raises the possibility that the ability of herbivores to successfully utilize plants might more readily evolve, while the ability of plants to evolve defenses will be more contingent (on local environments, etc.). This again supports a diffuse model of coevolution [45] and could eventually help us understand the accumulation of host-specific herbivores on plants through evolutionary time.

Genetic variation within species affects host-parasite interactions beyond herbivorous insects and their host plants [49], including for example susceptibility to parasitic diseases in humans and other animals being a function of both genetic variation in the hosts and among pathogen strains [50]. However, as is the case for plant-insect interactions, genomic investigations of other pairwise interactions have rarely considered both species simultaneously, but have focused on either the host or parasite. If epistatic, among-species interactions were common (as assumed by the gene-for-gene model of coevolution), the piecewise approach (focusing on one interacting species rather than the pair) might be a major roadblock to progress in understanding the evolution of these systems. However, if additivity and consistency of polygenic effects hold generally, as documented in the plant and herbivores studied here, a focus on one species in an interaction might not be misleading, and might inform predictive models, but this hypothesis remains to be tested with other interacting species.

## Methods

### Establishing the primary common garden

We planted a common garden comprising 1080 alfalfa plants at the Greenville Experimental Farm near Logan, Utah (41.765° N, 111.814° W) in 2018 (Fig. S1a). Seeds for this garden were collected from 11 naturalized alfalfa sites in the western USA, including five sites where *L. melissa* are found, and six sites lacking *L. melissa* butterflies (Table S1). An average of 4.9 seeds were planted from each of 220 maternal plants (with an average of 97.6 seeds planted from each site, SD = 8.6, range = 77 to 105).

We first germinated seeds in pots in a greenhouse in April 2018; the greenhouse was maintained at 24°C. Seeds were planted in Sun Gro Propagation mix # 3 (Sun Gro Horticulture, Agawam, MA, USA) after inoculation with Nitragin Gold Alfalfa (Monsanto, Creve Coeur, MO, USA) and scarification with sandpaper. Then, on May 24th 2018, we transplanted seedlings to the Greenville Experimental Farm on a 16×37 meter (m) plot. Plants were laid out in 15 rows of 72 plants each with 0.5 m spacing along rows and 1 m spacing between rows. We randomized plants with respect to source population and maternal family when planting. Plants were watered ~2–3 times per week using a sprinkler system. After the summer, most of the above ground biomass was removed from each plant, including any seed pods (this was done to prevent recruitment of additional plants). The garden was then allowed to overwinter (alfalfa is a perennial) before it was used the following summer (2019) for our experiment. During 2019, we again watered the garden ~2–3 times per week using a sprinkler system.

### Caterpillar husbandry and performance assays

We obtained *L. melissa* eggs from gravid females collected from six sites between June 16th and July 4th 2019 (Table S1). As in past work, gravid females were caged with a few sprigs of host plant (*M. sativa*) and allowed to lay eggs [16, 24, 26]. Eggs were kept in a Percival incubator (model no. 136VL) at 27°C with 14 hours light:10 hours dark. Upon hatching, caterpillars were assigned randomly to feed on a specific *M. sativa* plant. Each neonate caterpillar was carefully transferred to a Petri dish with a sprig of fresh plant material (a few leaflets) with the stem of the plant tissue wrapped in a damp Kimwipe. We verified each caterpillar was alive and uninjured after transfer. The Petri dish containing the caterpillar was then returned to the incubator. Caterpillars were given fresh leaf tissue *ad libitum* and were checked daily for survival, pupation and eclosion as adults.

As metrics of performance, we measured 8-day and 14-day caterpillar weight, and weight at pupation using a Mettler Toledo XPE105 analytical microbalance (Mettler Toledo). Weights were recorded to the nearest 0.01 mg, and we took the mean of two independent weight measurements. *Lycaeides melissa* caterpillars generally spend 20 to 30 days as larvae [16], and weight and lifetime fecundity are highly correlated in *L. melissa* [24]. We then considered the following nine performances metrics: 8-day caterpillar weight (mg), 14-day caterpillar weight (mg), weight at pupation (mg), survival to 8 days (binary), survival to 14 days (binary), survival to pupation (binary), survival to adult (binary), total survival time (integer valued), and truncated survival time (integer valued). For truncated survival time, we truncated survival at the maximum number of days required for any of the caterpillars to reach eclosion; this avoids caterpillars that developed slowly but never pupated or eclosed from having the longer survival times than caterpillar that successfully eclosed as adults.

### DNA extraction and sequencing

We isolated DNA from 1064 *M. sativa* plants from the Greenville Experimental Farm common garden (16 of the initial 1080 plants died before they could be used in the experiment) and 922 *L. melissa* caterpillars (some caterpillars were too small when they died to be recovered and used for DNA extraction), pupae or adults reared on these plants (DNA was also isolated from an additional 172 *M. sativa* and 157 *L. melissa* as part of a complementary smaller common garden experiment that is described below). *Medicago sativa* DNA was isolated from dried leaf tissue by Ag Biotech (Ag Biotech, Monterey CA, USA). We isolated DNA from whole caterpillars, pupa or the thorax of adult *L. melissa* butterflies using Qiagen’s DNeasy Blood & Tissue kit (Qiagen Inc. MA, USA) in accordance with the manufacturer’s recommendation.

We then used previously described procedures to create DNA-fragment libraries for our genotyping-by-sequencing approach [21, 51]. This approach has proven successful in the past for both *M. sativa* and *L. melissa* [21, 28, 52]. Briefly, we first digested the DNA from each sample with the restriction enzymes *EcoRI* and *MseI*. We then ligated adaptor oligonucleotides to the ends of the digested DNA fragments with *T4 DNA ligase*. The adaptor oligonulceotides included the Illumina adaptors and unique 8–10 base pair (bp) identification sequences or barcodes. Next, we PCR-amplified each fragment library as described in [21]. Amplified fragment libraries were then pooled (with sets of 96 samples combined), purified and size-selected. Size selection was accomplished using a BluePippin (Sage Science Inc., Beverly, MA, USA) by Utah State University’s Genomics Core Facility. Fragments between 300 and 450 bps in length were retained for both *M. sativa* and *L. melissa*. After purification and size selection, libraries were further pooled in groups of ~192 or ~384 samples for DNA sequencing.

We sequenced the DNA fragment libraries at the University of Texas Genomic Sequencing and Analysis Facility (Austin, TX, USA). Each of three pools of ~384 *M. sativa* and three pools of ~384 *L. melissa* were sequenced on one lane on a NovaSeq with S1 100 bp SR reads. A single pool of 184 *M. sativa* was sequenced on one lane of NovaSeq with SP 100 bp SR reads. After removing PhiX control sequences, this generated ~2.5 billion reads for *M. sativa* and ~2.5 billion reads for *L. melissa*.

### DNA sequence alignment and variant calling

We first de-multiplexed the *M. sativa* and *L. melissa* reads based on our internal barcode sequences; this was done using a custom `Perl` script. Next, we aligned the *M. sativa* sequences to the *M. sativa* genome [53]; this was done using the `mem` algorithm from `bwa` (version 0.7.17-r1188) [54]. For alignment, we considered internal seeds longer than 1.3× the minimum seed length (set to 15 bps) and only output alignments with a minimum mapping quality of 30. We then used `samtools` (version 1.10) to compress, sort and index the alignments [55]. Single nucleotide polymorphisms (SNPs) were next identified using `GATK` (version 4.1) [56]. We first used the `GATK HaplotypeCaller` to compute genotype likelihoods and generate `g.vcf` files. For this, we set the expected heterozygosity to 0.001 (across all sites), the minimum base quality to 20, and ploidy to four as *M. sativa* is a tetraploid species. We then used the `CombineGVCFs` and `GenotypeGVCFs` algorithms to call SNPs across the set of *M. sativa* individuals. An identical approach was used for generating and processing DNA sequence alignments for *L. melissa* except that sequences were aligned to the *L. melissa* genome [57, 58]. For *L. melissa*, we computed genotype likelihoods and identified SNPs using the consensus caller in `bcftools` (version 1.9) [55]. We first generated pileups with the `mpileup` algorithm using the recommended mapping quality adjustment (`-C 50`), a minimum mapping quality of 30, and a minimum base quality of 20. SNPs were then called using the `call` algorithm with a prior heterozygosity (`-P`) of 0.001 and a minimum posterior probability of a site being variable of 0.99.

We filtered the initial set of *M. sativa* SNPs to retain only SNPs with a mean sequence read depth (per individual) >2×, a minimum of 10 reads supporting the non-reference allele, a maximum absolute value for the base-quality rank-sum test of 3, a maximum absolute value for the mapping-quality rank-sum test of 3, a maximum absolute value for the read-position rank-sum test of 2.5, a minimum ratio of variant confidence to non-reference read depth of 2, a minimum mapping quality of 30, and missing data for no more than 20% of individuals. We further removed loci with >2 alleles or coverage exceeding 3 standard deviations above the mean. This left us with 161,008 SNPs. For *L. melissa*, we filtered the initial set of SNPs to retain only those with a mean sequence read depth (per individual) >2×, a minimum of 10 reads supporting the non-reference allele, a minimum *P*-value for a Mann-Whitney U test of base-quality bias of 0.01, a minimum *P*-value for a Mann-Whitney U test of read-position bias of 0.01, a minimum mapping quality of 30, missing data for no more than 20% of individuals, 2 alleles, and coverage not exceeding 3 standard deviations of the mean. This left us with 63,194 SNPs. Slight differences in filtering criteria for the two data sets reflect differences in the output provided by `GATK` (used for the tetraploid *M. sativa*) and `bcftools` (used for the diploid *L. melissa*).

### Inference of genotypes and genetic variation

We estimated genotypes using the Bayesian (ad)mixture model implemented in `entropy` (version 2.0) [21,59]. This program estimates genotypes for diploids or tetraploids while accounting for uncertainty caused by limited coverage and sequencing error, as captured by genotype likelihoods. The model assumes that the two (diploids) or four (tetraploids) allele copies at each SNP locus are drawn from unknown, hypothetical ancestral populations with each individual having a genome with ancestry from some mixture of these hypothetical populations. This allows the relevant allele frequencies for any given individual to be modeled as a mixture of allele frequency distributions. We estimated genotypes for *M. sativa* and *L. melissa* assuming two or three ancestral populations, and using the genotype likelihoods from `bcftools` (*L. melissa*) or `GATK` (*M. sativa*) as input. Estimates were obtained via Markov chain Monte Carlo (MCMC) with three chains, each with 10,000 iterations and a 5000 iteration burn-in. We set the thinning interval to 5. Point estimates of genotypes were obtained as the posterior mean estimate of the number of non-reference alleles, with the posterior summarized across chains and numbers of populations. Thus genotype estimates take on values between 0 and 2 for *L. melissa* and 0 and 4 for *M. sativa*, but are not constrained to be integer valued. We then visualized patterns of genetic variation using a principal component analysis (PCA) via the `prcomp` function in `R`, with the centered but not scaled genotype estimates as input (i.e., the covariance matrix). Genetic differentiation among experimental *M. sativa* or *L. melissa* based on their population of origin was characterized using Nei’s F_ST_ (also known as Nei’s G_ST_) [60,61]. Specifically, we estimated F_ST_ for each pair of plant or insect populations (localities) 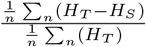.

### Preparing the caterpillar performance data for genetic mapping

We removed potential confounding variation from the caterpillar performance data prior to analyzing genotype-performance associations. First, we regressed each of the nine caterpillar performance metrics on caterpillar hatch date (to control for temporal effects) and source population. This was done with the `lm` function in `R`. Next, we used distance-based Moran’s eigenvector maps to remove possible effects of space (location) within the common garden. This procedure involves creating spatial variables based on a PCA of a truncated (nearest neighbors) Euclidean distance matrix (i.e., a principal coordinates analysis), where distance was defined from the spatial layout of the common garden [62]. We then used forward selection of variables following [63] to select spatial variables (eigenvectors) that explained the variation in each trait. Specifically, we first tested for a significant (at *P* < 0.05) fit of a model with all of the spatial variables. If and only if this full model was significant, we began adding spatial variables to a null model one at a time based on the extent to which they increased the total model *r*^2^. This procedure continued until either: (i) the *P*-value for the most recently added variable was > 0.05, (ii) the total *r*^2^ exceeded the original *r*^2^ from the full model with all variables, (iii) adding the new variable did not increase the model *r*^2^, or (iv) 200 spatial covariates had been added. The final models explained 18 to 51% of the variation in plant traits (mean = 35%) with 22 to 77 covariates retained; however, a model with no spatial covariates was selected for most caterpillar performance traits with 14-day weight being the sole exception (20 covariates explaining 14% of the trait variation). Scaled residuals from the final model for each trait were then used for genetic mapping.

### Multilocus genetic mapping of caterpillar performance

We tested for associations between (i) *M. sativa* SNPs (161,008 SNPs), (ii) *L. melissa* SNPs (63,194 SNPs), and (iii) SNPs from both species combined (224,202 SNPs), and each of the nine caterpillar performance metrics (i.e., the residuals from the models described in the previous paragraph). We performed these analyses using Bayesian sparse linear mixed models (BSLMMs), which we fit with `gemma` (version 0.95alpha) [64]. A key advantage of this approach for gentoype-phenotype association analyses is that, unlike traditional genome-wide association (GWA) mapping methods that test each genetic marker separately, the BSLMM approach fits all SNPs in a single model and thus mostly avoids issues related to testing large numbers of null hypotheses. The BSLMM method assumes that trait values are determined by a polygenic term and a vector of the (possible) measurable effects of each SNP on the trait (*β*) [64]. Bayesian Markov chain Monte Carlo (MCMC) with variable selection is used to infer the posterior inclusion probability (PIP) for each SNP, that is, the probability that each SNP has a non-zero effect or association, and the effect size conditional on it being non-zero [65]. The polygenic term denotes each individual’s expected deviation from the mean phenotype based on all of the SNPs. This term accounts for phenotypic covariances among individuals caused by their relatedness or overall genetic similarity [64]. The kinship matrix also serves to control for population structure and relatedness when estimating effects of individual SNPs (*β*) along with their PIPs. Similarly, SNPs in linkage disequilibrium (LD) with the same causal variant effectively account for each other, such that only one or the other is needed in the model, and this redundancy is captured by the posterior inclusion probabilities.

The hierarchical structure of the model makes it possible to estimate additional parameters that describe aspects of a trait’s genetic architecture [16, 52, 64, 65]. These include the percentage of the phenotypic variance explained (PVE) by additive genetic effects (which includes *β* and the polygenic term, and should approach the narrow-sense heritability), the percentage of the PVE due to SNPs with measurable effects or associations (PGE, the percentage of the phenotypic variance explained by genic effects, which is based only on *β*), and the number of SNPs with measurable associations (n-*γ*). All of these metrics use MCMC to integrate over uncertainty in the effects of individual SNPs, including whether these are non-zero. Lastly, using this BSLMM approach, it is also possible to obtain genomic-estimated breeding values (GEBVs) or polygenic scores, that is, the expected trait value for an individual from the additive effects of their genes, as captured by both *β* and the polygenic term [16, 52].

For each of the nine caterpillar performance metrics and three genetic data sets, we conducted 10 MCMC runs with `gemma`, each comprising 1 million iterations and a 200,000 iteration burn-in. Every 10th MCMC sample was retained to form the posterior distribution. Polygenic scores (i.e., genomic-estimated breeding values) were then calculated from the genetic data sets and model-averaged effect estimates for each SNP locus. Genetic covariance matrixes were computed from the estimated polygenic scores.

### Within-garden cross-validation and genomic prediction

We used 10-fold cross-validation to assess our ability to predict performance traits from *M. sativa* genetic data, *L. melissa* genetic data, and the combined genetic data from *M. sativa* and *L. melissa*. To do this, we first randomly assigned each observation to one of ten test data sets. Then, for each test data set, we estimated genotype-phenotype associations using `gemma` as described above, but based only on the 90% of individuals not in that test data set. For this, we used a single MCMC run comprising 1 million iterations, a 200,000 iteration burn-in, and a thinning interval of 10. We then used `gemma` to predict the phenotypes of the 10% of individuals held back for the test set (these individuals were not used to fit the model); this was done with the predict option in `gemma`. We then quantified predictive performance using the Pearson correlation between the genomic predictions of each performance metric and the observed values.

### Gene Miller Life Science Garden set up and genomic prediction

We further tested our ability to predict caterpillar performance trait values from genotypes by generating genomic predictions of performance for caterpillars reared on *M. sativa* from a second, smaller common garden comprising 180 *M. sativa*. This second garden, the Gene Miller Life Science Garden, planted on the Utah State University (USU) campus ~2.5 km from the main garden (41.742° N, 111.811° W), included plants from six of the 11 *M. sativa* source sites (30 plants per site from six maternal families) and caterpillars from each of the sites used in the main experiment (ALP, APLL, AWFS, BST, VUH, and VIC) (see Table S1 for site details). These plants were started in a greenhouse on April 18th 2018, with seeds sown in Sun Gro Propagation mix ≠ 3 (Sun Gro Horticulture, Agawam, MA, USA) after inoculation with Nitragin Gold Alfalfa (Monsanto, Creve Coeur, MO, USA) and scarification with sandpaper. The greenhouse was maintained at 24°C without overhead lights. The plants were watered ~3 times per week throughout the summer. Then, in the fall of 2018, we transplanted the 180 plants into the USU common garden. They were arranged in a randomized-block design with six blocks in the ~6 m × ~12 m garden.

We used leaf tissue from these plants for rearing *L. melissa* caterpillars in the summer of 2019 exactly as described for the main common garden at the Greenville Experimental Farm (see ‘Caterpillar husbandry and performance assays’ above for details). This parallel experiment was conducted at the same time as the main experiment. Plant and caterpillar samples from this parallel experiment were sequenced along with the samples from the Greenville Experimental Farm experiment. We successfully obtained genetic data from 172 *M. sativa* and 156 caterpillars of the 180 involved in this experiment. These genetic data were processed along with those from the main garden (see ‘DNA sequence alignment and variant calling’ above for details).

We then used the estimated, model-averaged effects from the BSLMM fits in `gemma` from the main garden to predict performance traits based on plant, caterpillar, or plant and caterpillar genotypes for these individuals. We compared these genomic predictions (i.e., polygenic scores computed from the main-garden models) to the observed performance trait values for these caterpillars. This was done using residuals after removing effects of hatch date and block (i.e., plot) within the USU garden. As with the within-garden cross-validation analyses described in the previous section, predictive power was measured by the Pearson correlation between the predicted and observed performance trait values.

### Plant trait measurements

We measured a series of morphological traits potentially associated with plant vigor or resistance to insects (e.g., putative structural plant defenses) [16, 66, 67] for each of the 1080 *M. sativa* plants in the Greenville Experimental Farm common garden. In May-June 2019, we measured leaf size (length, width and area), leaf shape (length/width), trichome density, dry leaf weight and specific leaf area (SLA) for each plant. Measurements were based on the middle leaflet from the terminal leaf taken from each of two haphazardly chosen sprigs from the center of the plant (taken from opposite sides). We measured the width (at the widest point) and length (along the midvein) of the middle leaflet with calipers (each leaf comprises three leaflets; measurements were taken to the nearest 1 mm). Next, we calculated leaf area (length × width) and shape (length/width) from these measurements. We then counted the number of trichomes in a ~2.5 mm diameter circle directly adjacent to the midvein under a stereoscope at 35× magnification. The two leaflets from each plant were then placed in a coin envelope in a bin with desiccant. The dry weight of these leaflets was measured on a Mettler Toledo XPE105 analytical microbalance (Mettler Toledo) to the nearest 0.01 mg. Leaf area and dry weight were used to calculate SLA (SLA is the ratio of leaf area to dry mass and is often correlated with leaf mechanical properties, such as work to tear, shear or punch) [67]. The other leaves from the two sprigs collected from each *M. sativa* plant were placed in coin envelopes in a bin with desiccant to be dried and preserved for chemical analyses (see the ‘Measuring plant chemistry’ in the next section). The height of each plant, measured from the top of the soil to the tip of the longest stem was also determined. Lastly, leaf toughness was measured using a penetrometer. Toughness was quantified as the mean force required to penetrate six leaflets at the midvein (three leaflets from each of two leaves, each from a different branch) (toughness was measured to the nearest 5 g).

In addition to these morphological traits, we quantified the extent to which each plant had been subject to herbivory in the field. This could be indicative both of the palatibility of the plant and of the extent to which it might be mounting an induced response to herbivory. Here, we recorded data from two branches on opposite sides of each plant. On each branch, we counted the number of fully open leaves from the fifth node to the terminal end of the branch, along with the number of these leaves with herbivore damage. From this, we computed the proportion of leaves with herbivory.

### Sample extraction and phytochemical analysis

Randomized chemical extractions of foliar tissues were carried out as previously described by [23]. Briefly, 10.0 mg of ground plant tissue (Quiagen Tissuelyser II, 30 Hz, 1 min) were combined with 2.00 mL of 70% aqueous ethanol and briefly vortexed before 15 minutes of sonication. Sample suspensions were then centrifuged at 500 rpm (Genevac EZ-2) for 10 min before filtering through 1 mL 96 well filter plates (glass fiber, 1 *μ*m, Pall, New York, NY) into 1 mL 96 well plates and sealed with a silicone plate mat (Agilent, Santa Clara, CA). Aliquots of 12 samples from each row were combined into pools. All samples (1 *μ*L) were co-injected with a 1 *μ*L air bubble and 1 *μ*L of digitoxin internal standard (ISD; 50 *μ*M in spectral grade MeOH, Fisher Optima). Between every two rows of samples, pools of those two rows were co-injected with ISD for retention time alignment, monitoring instrument response and structural determination. Analytical samples, pools and ISD blanks were injected onto an Agilent 1290 Infinity II UPLC equipped with a dual-channel variable wavelength detector (λ = 250 nm) connected to an Agilent 6560 ion-mobility-quadrupole-time-of-flight mass spectrometer equipped with a Jet Stream electrospray ionization dual source with reference mass infusion and tuned in 1700 m/z mode (IM-Q-TOF; drying gas temperature: 300 °C, drying gas flow: 5 L/m; drying gas flow: 8 L/m nebulizer pressure: 50 psig; sheath gas temperature: 275 °C; sheath gas flow: 8 L/m VCap: 3500 V; nozzle voltage: 1000 V; fragmentor: 300 V; octopole: 750 V). All standards, analytical samples and pools were analyzed in TOF mode (1 spectrum per second) and pools were also analyzed in iterative Auto Q-TOF mode (MS: 3 scans per second; MS/MS: 1 scan per second, collision energy: 20, 40 eV, precursor threshold: 10000 counts). Samples were eluted through an Agilent Poroshell 120 column (EC-C18, 1.9 μM, 2.1 × 100 mm) using a linear gradient comprised of solvent A (0.1% aqueous formic acid) and solvent B (99% acetonitrile containing 1% water and 0.1% formic acid) at 0.5 mL/min over 14 minutes as follows: 0 min: 5% B; 4 min: 50% B; 10-12 min: 100% B, ramp to 0.8 mL/min; 12.1-14 min: 1% B, ramp from 0.6 to 0.5 mL/min. Data were extracted and aligned using Agilent Profinder v10.0 before loading into Agilent Mass Profiler Professional v.15.1. Compounds in less than 10% of samples were excised before exporting peak areas to a csv file. Using the statistical platform R, peak areas were normalized to ISD and dry plant mass before statistical analysis. Tandem mass spectrometry (MS/MS) data were extracted using the find by Auto MS/MS function in Agilent Mass Hunter Qualitative Analysis before exporting to CEF files for further analysis.

### Multilocus genetic mapping of plant traits

We tested for associations between the *M. sativa* SNPs (161,008 SNPs) and 1760 plant traits: leaf length, leaf width, leaf area, leaf shape, leaf weight, SLA, trichome density, leaf toughness, plant height, field herbivory and 1750 metabolomic chemical features (see the previous two sections for details). This was done using the 1080 *M. sativa* plants from the main common garden at the Greenvile Experimental Farm in Logan, UT. We first removed possible effects of spatial location within the garden as captured by distance-based Moran’s eigenvector maps using forward selection of variables [63], exactly as described for the caterpillar performance traits above (see ‘Preparing the caterpillar performance data for genetic mapping’). As with the caterpillar performance traits, genotype-plant trait associations were estimated by fitting BSLMMs with `gemma` (version 0.95alpha) [64]. For each of the 1760 plant traits, we conducted 10 MCMC runs with `gemma`, each comprising 1 million iterations and a 200,000 iteration burn-in. Every 10th MCMC sample was retained to form the posterior distribution. Polygenic scores were then calculated from the genetic data sets and model-averaged effect estimates for each SNP locus. Genetic covariance matrixes were computed from the estimated polygenic scores. The model-fitting procedure was repeated with 1760 randomized plant trait data sets (i.e., values of each of the original traits were permuted among plants) to verify that the distribution genotype-phenotype associations from the real data set differed from null expectations.

### LASSO regression models

We used least absolute shrinkage and selection operator (LASSO) regression to (i) identify the subset of plant traits with polygenic scores that best predicted caterpillar-performance polygenic scores and (ii) estimate the direction and magnitude of these associations (as captured by the regression coefficients). This approach constitutes a form of regularized regression where a subset of regression coefficients are shrunk to zero [68]. Thus, in addition to inducing shrinkage on all coefficients, it serves as an approach for variable (feature) selection. We fit a LASSO regression model for the polygenic scores for each caterpillar performance trait with the 1760 plant trait polygenic scores as potential covariates. This was done with the `R` package `glmnet` (version 4.0-2) [69]. Ten-fold cross-validation was used to select a value for the penalty parameter λ. We then estimated the (shrunk) regression coefficients, the coefficient of determination (r^2^) and cross-validation *r*^2^ (squared correlation between observed and predicted values) using the optimal value for λ. An additional 10-fold cross-validation procedure was used to compute cross-validation *r*^2^ with the optimal value for λ. We then repeated this entire procedure after randomizing the caterpillar-performance polygenic scores for each performance trait (10 randomized data sets per caterpillar performance trait); this was done to gauge null expectations for the degree of variation in performance polygenic scores explained by chance associations with plant-trait polygenic scores.

We next asked whether plant-trait polygenic scores could explain and predict caterpillar performance at the phenotypic level (i.e., not just the polygenic scores). To do this, we fit analogous LASSO regression models to the observed performance trait data, or more specifically to the residuals from the observed data after removing the effects of hatch date and location in the common garden. Here too we analyzed randomized response variables as well to generate null expectations (100 randomized data sets for each performance trait).

Finally, we fit an additional set of LASSO regression models to evaluate the extent to which plant-genetic effects, as captured by the plant-trait polygenic scores, interacted with caterpillar genetics to affect performance. For this analysis, we summarized *L. melissa* caterpillar genetics using a PCA of the centered genotype matrix for the reared caterpillars. We used the first four PCs in the LASSO analysis; this allowed us to test for interactions without including a prohibitively large number covariates in the analysis. We specifically fit models for caterpillar-performance polygenic scores (as inferred from plant genetics) as a function of caterpillar genotype (PCs 1-4), plant-trait polygenic scores (1760 traits), and interactions between each plant-trait polygenic score and each of the four caterpillar genotype PCs. This was done with glmnet (version 4.0-2) [69] as described for the models without interactions.

### Structural annotations of phytochemicals

We annotated the 20 phytochemicals most strongly associated with caterpillar performance. Specifically, for each metabolite, we summed the absolute LASSO regression coefficients of association with the nine caterpillar performance polygenic scores, with each coefficient weighted by the heritability of the performance trait based on the plant genetic data, and then selected the 20 compounds with the largest sums for annotation. Three of these compounds were annotated to a high degree of certainty (Table S10) by comparison of their MS/MS spectra to experimental spectra. Compound PC.38.9 was identified as a phosphatidylcholine using Agilent Lipid Annotator v1.0, whereas tricin 7-glucoside and Apigenin 7-[p-coumaroyl-(→2)-[glucuronyl-(1→3)]-glucuronyl-(1→2)-glucuronide were identified based on similarity to experimental spectra found in HMDB [70] and to in silico generated MS/MS spectra using CFM-ID [71]. Lipids, peptides, and phosphatidylcholines (PC) were classified based on high-certainty chemical formulae generated by fragmentation trees created in the Sirius 4 platform for determination of structural information [72]. These formulae were either further classified by CANOPUS [73] within the Sirius 4 package, or by searching the formulae within the METLIN mass spectrometric database [74]. Compounds were classified as peptides if these were the only hits suggested in METLIN. One diglyceride was classified by chemical formula search in the LIPID MAPS structure database [75], and one PC was classified based on having similar fragmentation (20 eV; m/z =227.2002 [88%], 255.2319 [88%], 912.7012 [100%]) to another compound classified as a PC (MESA.122; 20 eV; m/z = 227.1999 [95%], 255.2315 [100%], 824.6479 [54%]), despite having different molecular masses. The remaining five compounds were not found during Auto MS/MS due to insufficient signal within the pools analyzed, although the precursor ions were observed. In these cases, one PC was classified by searching the LIPID MAPS structure database and four saponins were classified based on METLIN database hits. Of these, one has been previously found in *Medicago spp* [76]. (Medicagenic acid 3-O-beta-D-glucoside). Hits for Kudzusaponin SA2, Quillaic acid 3-[rhamnosyl-(1→3)-[galactosyl-(1→2)]-glucuronide], and 28-Glucosyloleanolic acid 3-[rhamnosyl-(1→2)-galactosyl-(1→3)-glucuronide] are likely glycosyl analogs of soyasapogeninol B, medicagenic or zanhic acids, respectively, which have been observed in *Medicago* [77]. While these structures this cannot be confirmed without MS/MS spectra, we are confident in their classification as saponins.

### Complementary USU greenhouse experiment

An additional rearing experiment was conducted to (i) replicate the general effect of *M. sativa* genotype on caterpillar performance and (ii) determine whether different plant genotypes had consistent effects of caterpillar performance across different butterfly populations and species. In this experiment, we did not analyze genotype directly, but instead used *M. sativa* source population as a proxy for genotype. Here, 1001 *M. sativa* plants from six source populations (ALP, APLL, AWFS, BST, VUH, and VIC) were grown from seed in a greenhouse at USU (Table S12). The plants were planted in the greenhouse on April 18th 2018, with seeds sown in Sun Gro Propagation mix ≠ 3 (Sun Gro Horticulture, Agawam, MA, USA) after inoculation with Nitragin Gold Alfalfa (Monsanto, Creve Coeur, MO, USA) and scarification with sandpaper. The greenhouse was maintained at 24°C without overhead lights. The plants were watered ~three times per week throughout the summer.

We reared at total of 672 caterpillars from four *L. melissa* populations (6-13 females per population; see Table S12), 133 *Colias eurytheme* caterpillars and 196 *Vanessa cardui* caterpillars on leaf tissue from the greenhouse-grown plants. *Lycaeides melissa* caterpillars were obtained from wild-caught gravid butterflies as described above (collections were made between June 4th and 20th 2018). *Colias eurytheme* is a legume specialist and caterpillars from this species were obtained by collecting gravid females from our *M. sativa* common garden at the Greenville Experimental Farm in Logan, UT (eggs were collected from 20 females on June 16th 2018). *Vanessa cardui* is a generalist butterfly that rarely feeds on alfalfa. We obtained caterpillars for this species by ordering eggs from Carolina Biological Supply Company (item number 144078, ordered five units of ~30-35 eggs May 22nd 2018) (Burlington, NC, USA).

Caterpillars were kept in incubators and fed plant material from the greenhouse-grown *M. sativa* in the same manner as for the main common garden at the Greenville Experimental Farm (see ‘Caterpillar husbandry and performance assays’ above for details). The only exception was that we ensured each caterpillar consumed leaves from a single source population (our proxy for genotype for this experiment) rather than a single individual. The specific plants used within each population were rotated haphazardly. As our metrics of caterpillar performance, we measured caterpillar weight at 8 and 14 days of development using a Mettler Toledo XPE105 analytical microbalance (Mettler Toledo). Survival to 8 days, 14 days, pupation and eclosion was also noted.

We then estimated the proportion of the variance in 8-day weight and 14-day weight partitioned among plant and caterpillar population by fitting linear mixed-effect models with restricted maximum likelihood. This was done using the `lmer` function from the `R` package `lme4` (version 1.1.23) (`R` version 4.0.2) [78]. We analyzed the data from the three butterfly species separately. Plant source population was included as a random effect for each analysis. Caterpillar population was included as a second random effect for *L. melissa* as the caterpillars came from four source populations. Caterpillar hatch data was included in the models for *L. melissa* and *C. eurytheme*, but not *V. cardui* as hatch occurred mostly on a single day. We evaluated the null hypothesis that the variance associated with each random effect was zero using an exact restricted likelihood ratio test [79, 80]. This was done with the function exactRLRT from the `R` package `RLRsim` (version 3.1.6) with the null distribution generated from 10,000 simulations [81].

To complement the mapping results above and provide an additional test of genetic variation in *M. sativa* for the morphological traits, we also measured plant height, leaf length, leaf width, leaf area, leaf shape, leaf weight, specific leaf are, leaf toughness and trichome density in the 2018 greenhouse experiment. Measurements were taken as described for the main garden during May 2018. We then estimated the proportion of trait variance attributed to plant population and family using linear mixed-effect models with restricted maximum likelihood. This was done using the `lmer` function from the `R` package `lme4` (version 1.1.23) (`R` version 4.0.2) [78]. We tested the null hypothesis that the variance associated with each random effect (population or family) was zero using an exact restricted likelihood ratio test [79, 80]. This was done with the function `exactRLRT` from the `R` package `RLRsim` (version 3.1.6) with the null distribution generated from 10,000 simulations [81].

### Complementary Nevada common garden rearing experiment

Three species of caterpillars were reared on alfalfa from an experimental garden at the University of Nevada, Reno (UNR Main Station; Fig. S1), previously described in [23]. Fifty-five individual plants were picked at random from the common garden and harvested throughout the summer of 2018 to support the growth of individually-reared caterpillars that were randomly assigned to specific plants (i.e., each caterpillar was fed the foliage from only one plant). One plant died during the summer, thus 54 plants were ultimately involved in analyses. Cuttings from plants were collected weekly and stored in a refrigerator before being fed to caterpillars in petri dishes (with leaves replenished every other day) in a growth chamber at 25C and a 12-hour light, 12-hour dark cycle. On day 10 of development, caterpillars were weighed to the nearest 0.01 mg on a Mettler Toledo XP26 microbalance as a measure of developmental performance.

The three caterpillar species used in experiments were *V. cardui*, *L. melissa*, and *C. eurytheme*. The first species (*V. cardui*) was ordered as eggs from Carolina Biological Supply Company; eggs of the second species (*L. melissa*) were obtained by caging wild-collected females from a location near Reno, Nevada; similarly, eggs of the third species came from females collected at an agricultural field in Northern California (39.1221°N, 121.9686° W). From each species, 10 caterpillars were assigned to each of the experimental plants (i.e., 550 caterpillars per species). For logistical reasons, we staggered the rearing with *V. cardui* starting on 29 June, *L. melissa* on 13 July, and *C. eurytheme* on 25 July, 2018.

For the results reported here, we calculated median weight per species per plant, and then used Pearson correlations between species (among individual plants) to ask if plants that support higher weight gain for one species of caterpillar support higher weight gain for other species.

## Acknowledgements

Thanks to Caitlyn Allgrunn, Jennifer Bryan, Hannah Curtis, Savannah Daines, Skylar Deschene, Shaylen Fidel, Daniel Johnson, Adair Schruhl, Amy Springer, Abbie Weight, and Roecale Yazzie for their help with planting and maintaining the common gardens and conducting the caterpillar rearing experiments. This work was funded by NSF awards to Z. Gompert (DEB 1638768), M. Forister (DEB 1638793), C. A. Buerkle (DEB 1638602), J. A. Fordyce and S. L. Lebeis (1638922) and C. C. Nice (1638773) and by a generous donation from Dr. Gene Miller that funded the Gene Miller Science Garden at USU.

## Author contributions

ZG, CAB, JAF, CCN, SLL, LKL, and MAF designed the study. ZG, TS, CP, SAY, EP, MES, JGH, CAB, LKL and MAF conducted the experiments and generated the data. ZG, TS, CP, and MAF analyzed the data. ZG, CP and MAF drafted the manuscript. All authors helped edit and revise the manuscript.

## Data availability

DNA sequence data have been archived on NCBIs SRA (Accession numbers pending). All other data are available from Dryad (DOI pending).

## Code availability

Computer code is available from `GitHub`, https://github.com/zgompert/DimensionsExperiment.

