## Supplemental Material for "Additive genetic effects in interacting species jointly determine the outcome of caterpillar herbivory"

### Supplemental Tables and Figures

Table S1: List of source sites for the *M. sativa* Greenville Experimental Farm common garden or for *L. melissa* caterpillars for the rearing experiment. Site abbreviations and full site names are given, along with whether each site was (Y=yes) or was not (N=no) a source for *L. melissa* and *M. sativa*

| Site | Site name | Latitude (°N) | Longitude (°W) | *L. melissa* | *M. sativa* |
|---|---|---|---|---|---|
| AFAL | Fallon, NV | 39.49 | 118.59 | N | Y |
| ALP | Alpine, WY | 43.17 | 111.01 | N | Y |
| APLL | Pole Line Road, CA | 38.59 | 121.73 | N | Y |
| AWFS | Patagonia, NV | 39.51 | 119.90 | N | Y |
| BST | Bonneville Shoreline Trail, UT | 41.73 | 111.79 | N | Y |
| CKV | Cokeville, WY | 42.00 | 110.94 | Y | Y |
| FRM | Mainstation Farm, NV | 39.51 | 119.72 | N | Y |
| FRP | Frary Peak, UT | 40.98 | 112.22 | Y | N |
| HBR | Heber, NV | 40.53 | 111.48 | N | Y |
| HLK | Honey Lake, CA | 40.24 | 120.31 | Y | N |
| LIK | Likely, CA | 41.23 | 120.50 | N | Y |
| SIN | Sinclair, WY | 41.86 | 107.09 | Y | N |
| VIC | Victor, ID | 43.66 | 111.11 | Y | Y |
| VUH | Verdi, NV | 39.51 | 112.00 | Y | Y |

Table S2: Bayesian estimates of the proportion of genetic variation in caterpillar performance traits explained by *M. sativa* genetics, *L. melissa* genetics, or both combined as inferred from fitting Bayesian sparse linear mixed models with `gemma`. Traits shown are W8d = 8-day weight, W14d = 14-day weight, Wpup = pupal weight, S8d = 8-day survival, S14d = 14-day survival, SPup = survival to pupation, SAdu = survival to adult, Stot = total survival time, and Stime = (truncated) survival time. Posterior medians (med) and lowers (lb) and upper (ub) of Bayesian equal-tail probability intervals are given.

| | *M. sativa* | | | *L. melissa* | | | Combined | | |
|---|---|---|---|---|---|---|---|---|---|
| Trait | med | lb | ub | med | lb | ub | med | lb | ub |
| W8d | 0.11 | 0.03 | 0.23 | 0.29 | 0.20 | 0.38 | 0.32 | 0.17 | 0.48 |
| W14d | 0.36 | 0.23 | 0.50 | 0.12 | 0.04 | 0.23 | 0.49 | 0.34 | 0.64 |
| WPup | 0.16 | 0.02 | 0.42 | 0.05 | 0.00 | 0.19 | 0.23 | 0.03 | 0.53 |
| S8d | 0.02 | 0.00 | 0.09 | 0.21 | 0.14 | 0.27 | 0.17 | 0.10 | 0.25 |
| S14d | 0.03 | 0.00 | 0.10 | 0.19 | 0.13 | 0.25 | 0.16 | 0.11 | 0.22 |
| SPup | 0.21 | 0.12 | 0.31 | 0.05 | 0.01 | 0.13 | 0.26 | 0.14 | 0.38 |
| SAdu | 0.13 | 0.04 | 0.24 | 0.11 | 0.04 | 0.21 | 0.17 | 0.07 | 0.28 |
| Stot | 0.04 | 0.00 | 0.11 | 0.20 | 0.13 | 0.26 | 0.17 | 0.10 | 0.26 |
| Stime | 0.10 | 0.02 | 0.20 | 0.14 | 0.08 | 0.20 | 0.18 | 0.10 | 0.30 |

Table S3: Bayesian estimates of the proportion of genetic variation in caterpillar performance traits explained by *M. sativa* genetics, *L. melissa* genetics, or both combined that is explained by measurable SNP-performance associations as inferred from fitting Bayesian sparse linear mixed models with `gemma`. Traits shown are W8d = 8-day weight, W14d = 14-day weight, Wpup = pupal weight, S8d = 8-day survival, S14d = 14-day survival, SPup = survival to pupation, SAdu = survival to adult, Stot = total survival time, and Stime = (truncated) survival time. Posterior medians (med) and lowers (lb) and upper (ub) of Bayesian equal-tail probability intervals are given.

| | *M. sativa* | | | *L. melissa* | | | Combined | | |
|---|---|---|---|---|---|---|---|---|---|
| Trait | med | lb | ub | med | lb | ub | med | lb | ub |
| W8d | 0.31 | 0.00 | 0.91 | 0.84 | 0.61 | 0.98 | 0.35 | 0.09 | 0.78 |
| W14d | 0.30 | 0.00 | 0.87 | 0.28 | 0.00 | 0.85 | 0.24 | 0.01 | 0.73 |
| WPup | 0.26 | 0.00 | 0.89 | 0.34 | 0.00 | 0.93 | 0.25 | 0.00 | 0.88 |
| S8d | 0.35 | 0.00 | 0.93 | 0.97 | 0.89 | 1.00 | 0.90 | 0.68 | 0.99 |
| S14d | 0.34 | 0.00 | 0.93 | 0.95 | 0.83 | 1.00 | 0.89 | 0.66 | 0.99 |
| SPup | 0.24 | 0.02 | 0.63 | 0.30 | 0.00 | 0.90 | 0.12 | 0.00 | 0.55 |
| SAdu | 0.17 | 0.00 | 0.78 | 0.72 | 0.16 | 0.98 | 0.43 | 0.11 | 0.86 |
| Stot | 0.44 | 0.00 | 0.94 | 0.88 | 0.68 | 0.99 | 0.81 | 0.54 | 0.98 |
| Stime | 0.18 | 0.00 | 0.82 | 0.68 | 0.41 | 0.94 | 0.46 | 0.25 | 0.84 |

Table S4: Bayesian estimates of the number of genetic variants with measurable associations (effects) on caterpillar performance in *M. sativa* genetics, *L. melissa* genetics, or both combined as inferred from fitting Bayesian sparse linear mixed models with `gemma`. Traits shown are W8d = 8-day weight, W14d = 14-day weight, Wpup = pupal weight, S8d = 8-day survival, S14d = 14-day survival, SPup = survival to pupation, SAdu = survival to adult, Stot = total survival time, and Stime = (truncated) survival time. Posterior medians (med) and lowers (lb) and upper (ub) of Bayesian equal-tail probability intervals are given.

| | *M. sativa* | | | *L. melissa* | | | Combined | | |
|---|---|---|---|---|---|---|---|---|---|
| Trait | med | lb | ub | med | lb | ub | med | lb | ub |
| W8d | 23 | 0 | 211 | 13 | 7 | 23 | 7 | 1 | 206 |
| W14d | 100 | 1 | 286 | 8 | 0 | 161 | 51 | 1 | 263 |
| WPup | 12 | 0 | 189 | 10 | 0 | 178 | 15 | 0 | 160 |
| S8d | 6 | 0 | 240 | 10 | 5 | 16 | 5 | 2 | 10 |
| S14d | 5 | 0 | 128 | 6 | 3 | 12 | 3 | 2 | 5 |
| SPup | 8 | 1 | 75 | 14 | 0 | 247 | 24 | 0 | 233 |
| SAdu | 12 | 0 | 237 | 7 | 2 | 31 | 5 | 1 | 70 |
| Stot | 4 | 0 | 55 | 5 | 3 | 11 | 3 | 2 | 7 |
| Stime | 21 | 0 | 246 | 1 | 1 | 4 | 1 | 1 | 2 |

Table S5: Summary of traits and genes associated with SNPs with posterior inclusion probabilities (PIPs) of $> 0.5$ for at least on caterpillar performance trait. The chromosome and position of each SNP is given, along with the trait(s) for which it has a PIP $> 0.5$. Traits shown are W8d = 8-day weight, S8d = 8-day survival, S14d = 14-day survival, Stot = total survival time, and Stime = (truncated) survival time. Then, for each SNP, the nearest annotated gene (if <50 kbps away) is given, along with the distance to the boundary of the gene. No chromosome is given for the last SNP, but instead the scaffold number if reported (1260); this scaffold most likely corresponds to the genome of the bacterial endosymbiont *Wolbachia*.

| Chromosome | Position (bp) | Traits | Gene ID | Distance (kbp) |
|---|---|---|---|---|
| 1 | 18,994,990 | S8d | NA | NA |
| 1 | 26,801,442 | S8d, S14d, Stot | NA | NA |
| 2 | 5,400,668 | W8d | *Vacuolar protein sorting-associated protein 13* | 4 |
| 2 | 8,585,087 | Sadu | *V-type proton ATPase 116 kDa subunit a* | 13 |
| 5 | 14,087,514 | Stot | NA | NA |
| 8 | 10,131,457 | W8d | *Nesprin-1/MSP-300* | 2 |
| 8 | 14,984,485 | W8d | *Lipase member H* | <1 |
| 8 | 15,768,108 | W8d | *Juvenile hormone acid O-methyltransferase* | 9 |
| 16 | 10,535,866 | W8d | NA | NA |
| NA-1260 | 1,620,413 | Stime | *Ankyrin* | 8 |

Table S6: Summary of *M. sativa* genes within 30 kbps of the single SNP (chromosome 1, position 12,930,966) strongly associated with caterpillar performance (posterior inclusion probability for survival to pupation = 0.65). Gene locations (boundaries) were taken from the *M. sativa* genome annotation [53], and gene identifications were then obtained by submitting the gene sequence to NCBI BLAST data base via the megablast algorithm (accessed December 13th, 2021).

| Gene ID | Location (bps) | Distance to SNP (bps) |
|---|---|---|
| *Dentin sialophosphoprotein* | 12,898,211–12,909,201 | 21,765 |
| Unknown gene | 12,913,180–12,915,634 | 15,332 |
| *Photosystem I reaction center subunit psaK* | 12,917,170–12,918,098 | 12,868 |
| *DEAD-box ATP-dependent RNA helicase 35* | 12,924,807–12,926,582 | 4384 |
| *TOM1-like protein 9* | 12,928,300–12,937,724 | 0 |
| *D-amino-acid transaminase* | 12,946,628–12,949,340 | 15,662 |
| *Transcription factor MYB3R-1* | 12,951,386–12,957,121 | 20,420 |

Table S7: Summary of genetic contributions to plant trait variation in the main Greenville Farm common garden and 2018 greenhouse experiment. PVE denotes the proportion of trait variation explained by genetic effects from the Bayesian multilocus genetic mapping model applied to the common garden experiment; posterior medians (med) and lower (lb) and upper bounds (ub) of the 95% equal-tail probability interval are given. Variance components (var) for the contribution of plant population and plant family to the trait variation in the greenhouse experiment are also given, along with restricted likelihood ratio test statistic (RLRT) and associated $P$ values for the null that the variance component is 0. Results are shown for leaf length, leaf width, leaf area, leaf weight, specific leaf area (SLA), trichome density, plant height, leaf toughness and field herbivory levels (common garden only).

| | PVE | | | Plant pop. | | | Plant fam. | | |
|---|---|---|---|---|---|---|---|---|---|
| Trait | med | lb | ub | var | RLRT | $P$ | var | RLRT | $P$ |
| Leaf length | 0.09 | 0.02 | 0.19 | 0.06 | 21.88 | <0.001 | 0.18 | 38.47 | <0.001 |
| Leaf width | 0.39 | 0.28 | 0.49 | 0.09 | 32.63 | <0.001 | 0.27 | 76.36 | <0.001 |
| Leaf area | 0.09 | 0.01 | 0.20 | 0.10 | 37.69 | <0.001 | 0.25 | 65.39 | <0.001 |
| Leaf shape | 0.28 | 0.18 | 0.39 | 0.01 | 1.52 | 0.076 | 0.17 | 34.53 | <0.001 |
| Leaf weight | 0.34 | 0.21 | 0.46 | 0.06 | 20.92 | <0.001 | 0.23 | 58.50 | <0.001 |
| SLA | 0.12 | 0.02 | 0.23 | 0.04 | 12.08 | <0.001 | 0.13 | 12.93 | <0.001 |
| Trichomes | 0.20 | 0.10 | 0.31 | 0.02 | 2.85 | 0.033 | 0.18 | 36.56 | <0.001 |
| Height | 0.28 | 0.19 | 0.36 | 0.10 | 38.24 | <0.001 | 0.37 | 130.74 | <0.001 |
| Toughness | 0.20 | 0.10 | 0.31 | 0.00 | 0.06 | 0.326 | 0.09 | 10.68 | <0.001 |
| Herbivory | 0.06 | 0.01 | 0.15 | NA | NA | NA | NA | NA | NA |

Table S8: Variance in caterpillar performance explained by plant trait polygenic scores. Results are shown for lasso regression models of caterpillar performance polygenic scores inferred from *M. sativa* genetics and for the observed phenotypes. Variance explained is measured by $r^2$, whereas predictive variance explained is measured by cross-validation (CV) $r^2$. Caterpillar performance traits shown are W8d = 8-day weight, W14d = 14-day weight, Wpup = pupal weight, S8d = 8-day survival, S14d = 14-day survival, SPup = survival to pupation, SAdu = survival to adult, Stot = total survival time, and Stime = (truncated) survival time.

| | Poly. score | | Obs. phenotype | |
|---|---|---|---|---|
| Trait | $r^2$ | CV $r^2$ | $r^2$ | CV $r^2$ |
| W8d | 0.80 | 0.76 | 0.12 | 0.02 |
| W14d | 0.61 | 0.58 | 0.07 | 0.04 |
| wPup | 0.71 | 0.68 | 0.10 | 0.01 |
| S8d | 0.76 | 0.73 | 0.00 | 0.00 |
| S14d | 0.76 | 0.73 | 0.00 | 0.00 |
| SPup | 0.72 | 0.68 | 0.09 | 0.05 |
| SAdu | 0.71 | 0.66 | 0.06 | 0.00 |
| Stot | 0.41 | 0.39 | 0.00 | 0.00 |
| Stime | 0.75 | 0.71 | 0.00 | 0.00 |

Table S9: Variance in caterpillar performance explained by plant trait polygenic scores and interactions with caterpillar genetics (PCs 1-4). Results are shown for lasso regression models of caterpillar performance polygenic scores inferred from *M. sativa* genetics and for the observed phenotypes. Variance explained is measured by $r^2$, whereas predictive variance explained is measured by cross-validation (CV) $r^2$. Caterpillar performance traits shown are W8d = 8-day weight, W14d = 14-day weight, Wpup = pupal weight, S8d = 8-day survival, S14d = 14-day survival, SPup = survival to pupation, SAdu = survival to adult, Stot = total survival time, and Stime = (truncated) survival time.

| Trait | $r^2$ | CV $r^2$ |
|---|---|---|
| W8d | 0.62 | 0.48 |
| W14d | 0.40 | 0.25 |
| WPup | 0.48 | 0.32 |
| S8d | 0.55 | 0.29 |
| S14d | 0.52 | 0.26 |
| SPup | 0.62 | 0.27 |
| SAdu | 0.85 | 0.28 |
| Stot | 0.34 | 0.17 |
| Stime | 0.64 | 0.32 |

Table S10: Annotation of the 20 chemicals most strongly associated with polygenic scores for caterpillar performance. Details on columns from Casey XXX.

| ID | RT | Mass | MS/MS | Formula | Class | Class Source |
|---|---|---|---|---|---|---|
| Apigenin 7-[p-coumaroyl-(→2)-[glucuronyl-(1→3)]-glucuronyl-(1→2)-glucuronide] | 2.727 | 944.1856 | Y | $C_{42}H_{40}O_{25}$ | Saponin | CFM-ID |
| Medicagenic acid 3-O-beta-D-glucoside (MESA.1112) | 3.892 | 664.3815 | N | $C_{36}H_{56}O_{11}$ | Saponin | METLIN |
| Peptide (MESA.1185) | 2.647 | 472.196 | Y | $C_{18}H_{28}N_6O_9$ | Peptide | METLIN |
| PC(P-18:1(9Z)/22:2(13Z,16Z)) (MESA.122) | 12.5 | 823.6381 | Y | $C_{48}H_{90}NO_7P$ | Phosphatidyl Choline | CANOPUS |
| Diglyceride (MESA.124) | 11.75 | 712.5103 | Y | $C_{47}H_{68}O_5$ | Lipids | Lipid Maps |
| 28-Glucosyloleanolic acid 3-[rhamnosyl-(1→2)-galactosyl-(1→3)-glucuronide] (MESA.1305) | 4.212 | 1116.5345 | N | $C_{54}H_{84}O_{24}$ | Saponin | METLIN |
| PC (MESA.1342) | 9.146 | 803.532 | N | $C_{46}H_{78}NO_8P$ | Phosphatidyl Choline | Lipid Maps |
| Tricin 7-glucoside (MESA.143) | 3.317 | 492.1274 | Y | $C_{23}H_{24}O_{12}$ | Flavonoid Glycosides | CFM-ID |
| PC (MESA.339) | 12.35 | 911.6881 | Y | $C_{48}H_{98}NO_{12}P$ | Phosphatidyl Choline | Lipid maps |
| Fatty Acid (MESA.438) | 5.138 | 290.1886 | Y | $C_{18}H_{26}O_3$ | Lipids | CANOPUS |
| Peptide (MESA.545) | 2.085 | 407.1688 | Y | $C_{20}H_{26}N_2O_7$ | Peptide | CANOPUS |
| Fragment of MESA.615 (MESA.583) | 3.182 | 190.1359 | Y | $C_{13}H_{18}$ | Peptide | |
| Peptide (MESA.584) | 2.512 | 472.1963 | Y | $C_{18}H_{28}N_6O_9$ | Peptide | METLIN |
| Peptide (MESA.615) | 3.204 | 529.3227 | Y | $C_{23}H_{43}N_7O_7$ | Peptide | CANOPUS |
| Kudzusaponin SA2 (MESA.730) | 3.367 | 944.4944 | N | $C_{47}H_{76}O_{19}$ | Saponin | METLIN |
| Peptide (MESA.784) | 2.977 | 392.1101 | Y | $C_{15}H_{16}N_6O_7$ | Peptide | CANOPUS |
| Peptide (MESA.849) | 3.925 | 249.1336 | Y | $C_{14}H_{19}NO_3$ | Peptide | CANOPUS |
| Peptide (MESA.972) | 2.19 | 537.2772 | Y | $C_{30}H_{39}N_3O_6$ | Peptide | METLIN |
| PC 16:3_22:6 | 9.94 | 799.5293 | Y | $C_{46}H_{74}NO_8P$ | Phosphatidyl Choline | LipidAnnotator |
| Quillaic acid 3-[rhamnosyl-(1→3)-[galactosyl-(1→2)]-glucuronide] | 4.262 | 970.4766 | N | $C_{48}H_{74}O_{20}$ | Saponin | METLIN |

Table S11: Summary of associations of plant chemicals with caterpillar performance traits based on plant polygenic scores. Standardized LASSO regression coefficients are shown for the 20 phytochemicals most strongly associated with performance across the 9 performance traits. See Table S10 for full details of chemical identifications. The caterpillar performance traits are W8d = 8-day weight, W14d = 14-day weight, Wpup = pupal weight, S8d = 8-day survival, S14d = 14-day survival, SPup = survival to pupation, SAdu = survival to adult, Stot = total survival time, and Stime = (truncated) survival time.

| Chemical | W8d | W14d | WPup | S8d | S14d | SPup | SAdu | Stot | Stime |
|---|---|---|---|---|---|---|---|---|---|
| Apigenin | 0.00 | 0.00 | 0.00 | 0.00 | 0.00 | 0.05 | 0.03 | 0.00 | 0.01 |
| MESA.1112 | 0.00 | 0.00 | 0.00 | 0.04 | 0.04 | 0.03 | 0.02 | 0.04 | 0.06 |
| MESA.1185 | 0.00 | 0.00 | 0.00 | -0.06 | -0.06 | -0.04 | 0.00 | -0.03 | -0.06 |
| MESA.122 | 0.00 | -0.05 | 0.00 | 0.00 | 0.00 | 0.00 | 0.00 | 0.00 | 0.00 |
| MESA.124 | 0.00 | 0.00 | 0.00 | 0.00 | 0.00 | 0.09 | 0.06 | 0.00 | 0.07 |
| MESA.1305 | 0.11 | 0.00 | 0.00 | 0.00 | 0.00 | 0.00 | 0.00 | 0.00 | 0.00 |
| MESA.1342 | 0.11 | 0.00 | 0.00 | 0.00 | 0.00 | 0.00 | 0.07 | 0.00 | 0.01 |
| MESA.143 | 0.00 | 0.00 | 0.00 | 0.05 | 0.03 | 0.01 | 0.03 | 0.00 | 0.07 |
| MESA.339 | 0.00 | 0.03 | 0.10 | 0.00 | -0.03 | 0.00 | 0.00 | 0.00 | 0.00 |
| MESA.438 | 0.00 | 0.00 | -0.00 | 0.00 | -0.00 | -0.00 | -0.11 | 0.00 | -0.08 |
| MESA.545 | -0.03 | -0.02 | 0.00 | 0.00 | 0.00 | -0.01 | 0.00 | 0.00 | 0.00 |
| MESA.583 | 0.02 | 0.00 | 0.00 | -0.02 | -0.06 | -0.06 | -0.02 | -0.05 | -0.08 |
| MESA.584 | 0.00 | -0.09 | 0.00 | 0.00 | 0.00 | 0.00 | 0.00 | 0.00 | 0.00 |
| MESA.615 | 0.00 | 0.00 | -0.02 | 0.00 | 0.00 | -0.03 | -0.04 | 0.00 | -0.02 |
| MESA.730 | 0.01 | 0.02 | 0.01 | 0.04 | 0.02 | 0.04 | 0.02 | 0.00 | 0.03 |
| MESA.784 | -0.02 | -0.03 | -0.01 | 0.00 | 0.00 | 0.00 | -0.01 | 0.00 | 0.00 |
| MESA.849 | -0.08 | -0.05 | 0.00 | 0.00 | 0.00 | -0.00 | -0.05 | 0.00 | 0.00 |
| MESA.972 | 0.01 | 0.01 | 0.01 | 0.00 | 0.00 | 0.04 | 0.03 | 0.00 | 0.00 |
| PC 16:3_22:6 | 0.00 | 0.04 | 0.00 | 0.00 | 0.00 | 0.05 | 0.00 | 0.00 | 0.00 |
| Quillaic acid | -0.02 | -0.02 | 0.00 | 0.00 | 0.00 | 0.00 | 0.00 | 0.00 | 0.00 |

Table S12: List of source sites for the *M. sativa* or *L. melissa* caterpillars for the 2018 greenhouse experiment at Utah State University. Site abbreviations and full site names are given, along with sample sizes for *L. melissa* and *M. sativa*. 133 *Colias eurytheme* caterpillars were obtained from the Greenville Experimental Farm in Logan, UT; 196 *Vanessa cardui* were obtained from an online supplier, Carolina Biological Supply.

| Site | Site name | Latitude (°N) | Longitude (°W) | No. *L. melissa* | No. *M. sativa* |
|---|---|---|---|---|---|
| ALP | Alpine, WY | 43.17 | 111.01 | NA | 167 |
| APLL | Pole Line Road, CA | 38.59 | 121.73 | NA | 171 |
| AWFS | Patagonia, NV | 39.51 | 119.90 | NA | 162 |
| BST | Bonneville Shoreline Trail, UT | 41.73 | 111.79 | 191 | 166 |
| BWP | Beckworth Pass, CA | 39.78 | 120.07 | 192 | NA |
| HWR | Hardware Ranch, UT | 41.61 | 111.62 | 99 | NA |
| JJT | Jardine Juniper Trail, UT | 41.80 | 111.65 | 190 | NA |
| VIC | Victor, ID | 43.66 | 111.11 | NA | 170 |
| VUH | Verdi, NV | 39.51 | 112.00 | NA | 165 |

Table S13: Variance components describing the contribution of plant or caterpillar population to caterpillar performance for the 2018 USU greenhouse rearing experiment,along with restricted likelihood ratio test statistic (RLRT) and associated $P$ values for the null that the variance component is 0.

| | | Plant pop. | | | Insect pop. | | |
|---|---|---|---|---|---|---|---|
| Species | Trait | % var. | RLRT | $P$ | % var. | RLRT | $P$ |
| *L. melissa* | 8d wgt. | 7.96% | 16.13 | $< 0.001$ | 3.99% | 14.09 | $< 0.001$ |
| | 14d wgt. | 14.1% | 56.88 | $< 0.001$ | $\sim$0.00% | 0 | 1.0 |
| *C. eurytheme* | 8d wgt. | 2.97% | 0.70 | 0.164 | NA | NA | NA |
| | 14d wgt. | 9.40% | 4.29 | 0.014 | NA | NA | NA |
| *V. cardui* | 8d wgt. | 10.1% | 3.44 | 0.023 | NA | NA | NA |
| | 14d wgt. | 13.0% | 4.43 | 0.012 | NA | NA | NA |

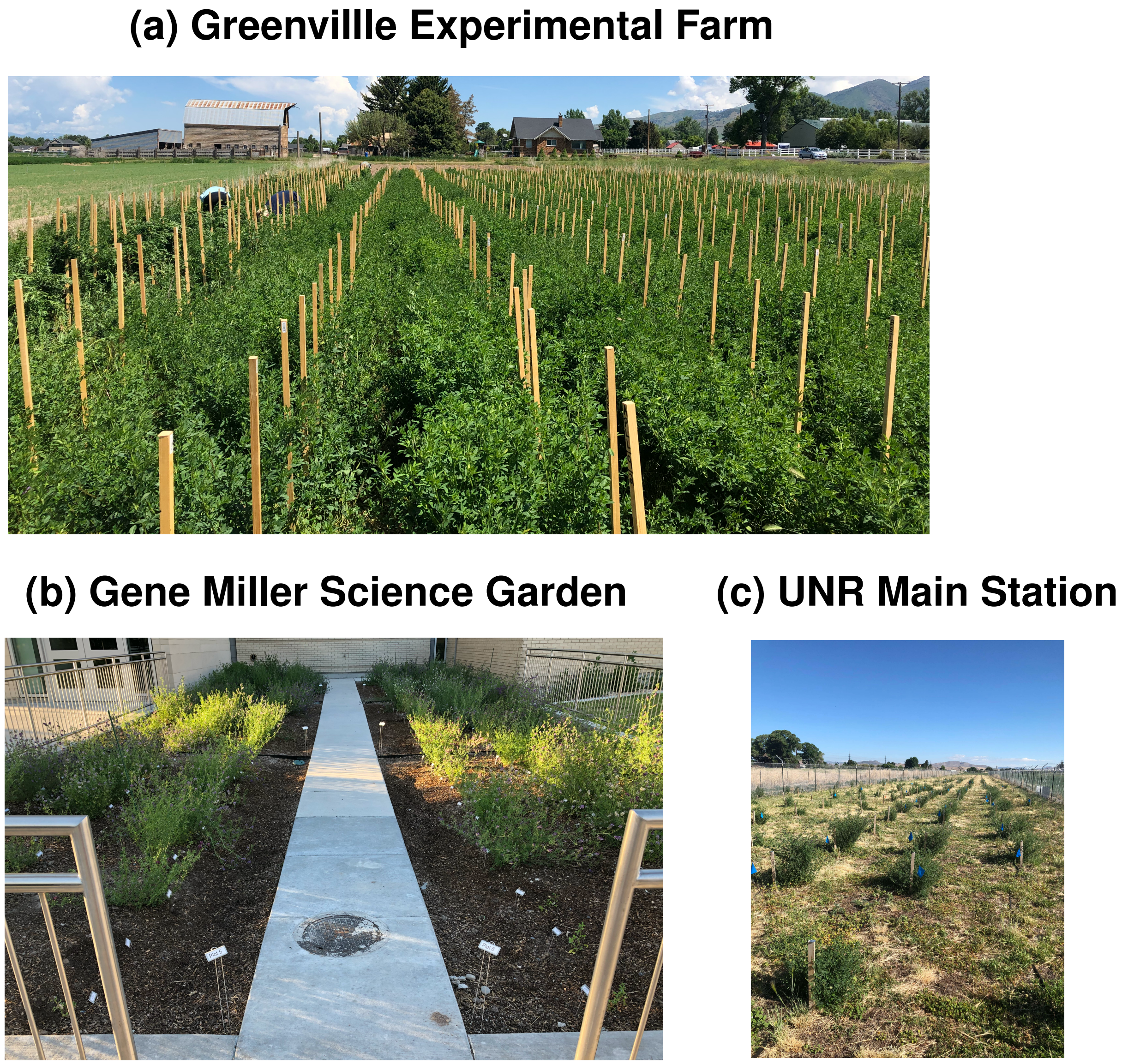


Figure S1: Photographs of the Greenville Experimental Farm in Logan, UT (i.e., the main common garden) (a) the Gene Miller Science Garden in Logan, UT (b) and the UNR Main Station garden in Reno, NV (c).

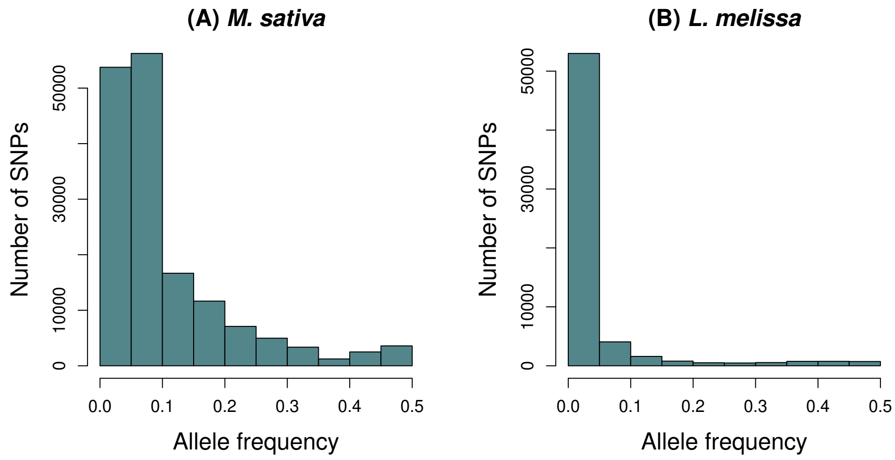


Figure S2: Histograms show the minor allele frequency distributions for SNPs in the Greenville Experimental Farm (i.e., the main common garden) *M. sativa* (a) and *L. melissa* caterpillars reared on these plants (b).

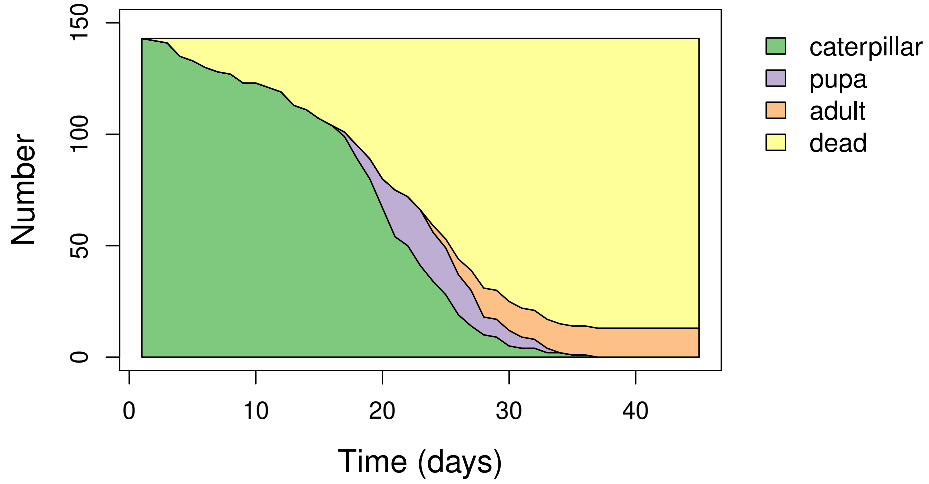


Figure S3: Plot shows survival and development of *L. melissa* over the course of the rearing experiment for caterpillars fed plants from the test science garden. Colored regions denote the number of individuals that were living caterpillar, pupa, adults or dead at each day post hatching.

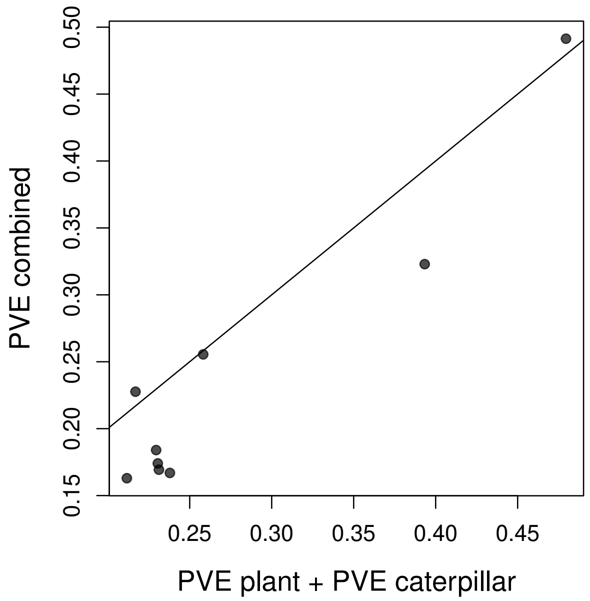


Figure S4: The scatterplot shows estimates of the proportion of variance explained (PVE) for each of nine caterpillar performance traits based on models of caterpillar and plant genetics fit separately with the PVEs added (x-axis) versus a single model fit with both caterpillar and plant genetics (y-axis). A 1:1 line is shown for reference.

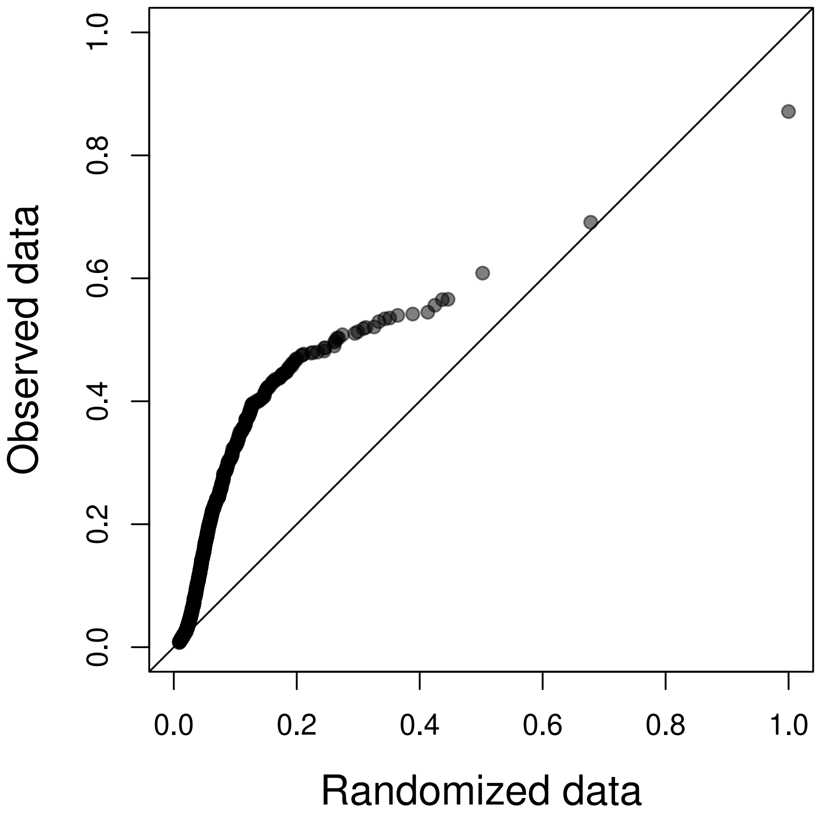


Figure S5: Quantile-quantile plot showing estimates of the proportion of variation explained (PVE) for each of 1750 plant chemistry traits (y-axis) versus the PVE for 1750 randomized variables each obtained by permuting on of the 1750 chemistry traits with respect to plant genetic data. A 1:1 line is shown for reference.

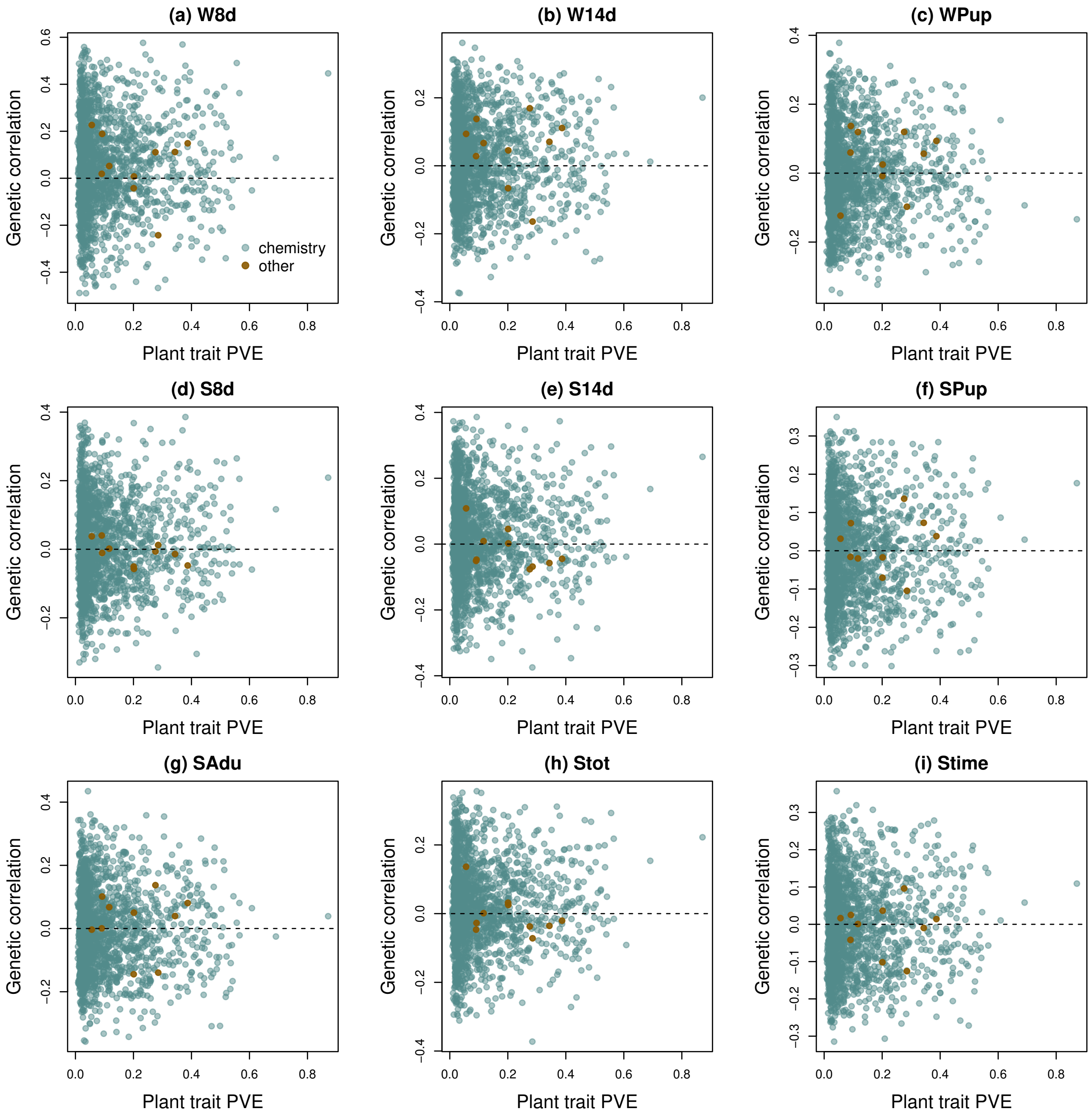


Figure S6: Scatterplots show the proportion of variance explained (PVE) by plant genetics for each of 1760 plant traits in relation to the genetic correlation between each plant trait and each of nine caterpillar performance traits. Polygenic scores for the performance traits were estimates solely from the plant genetic data. Caterpillar performance traits shown are W8d = 8-day weight, W14d = 14-day weight, Wpup = pupal weight, S8d = 8-day survival, S14d = 14-day survival, SPup = survival to pupation, SAdu = survival to adult, Stot = total survival time, and Stime = (truncated) survival time. Points are colored to reflect whether they are plant chemistry traits (blue, 1750 traits) or other plant traits (brown, 10 traits). A dashed line separates positive and negative genetic correlations in each panel.

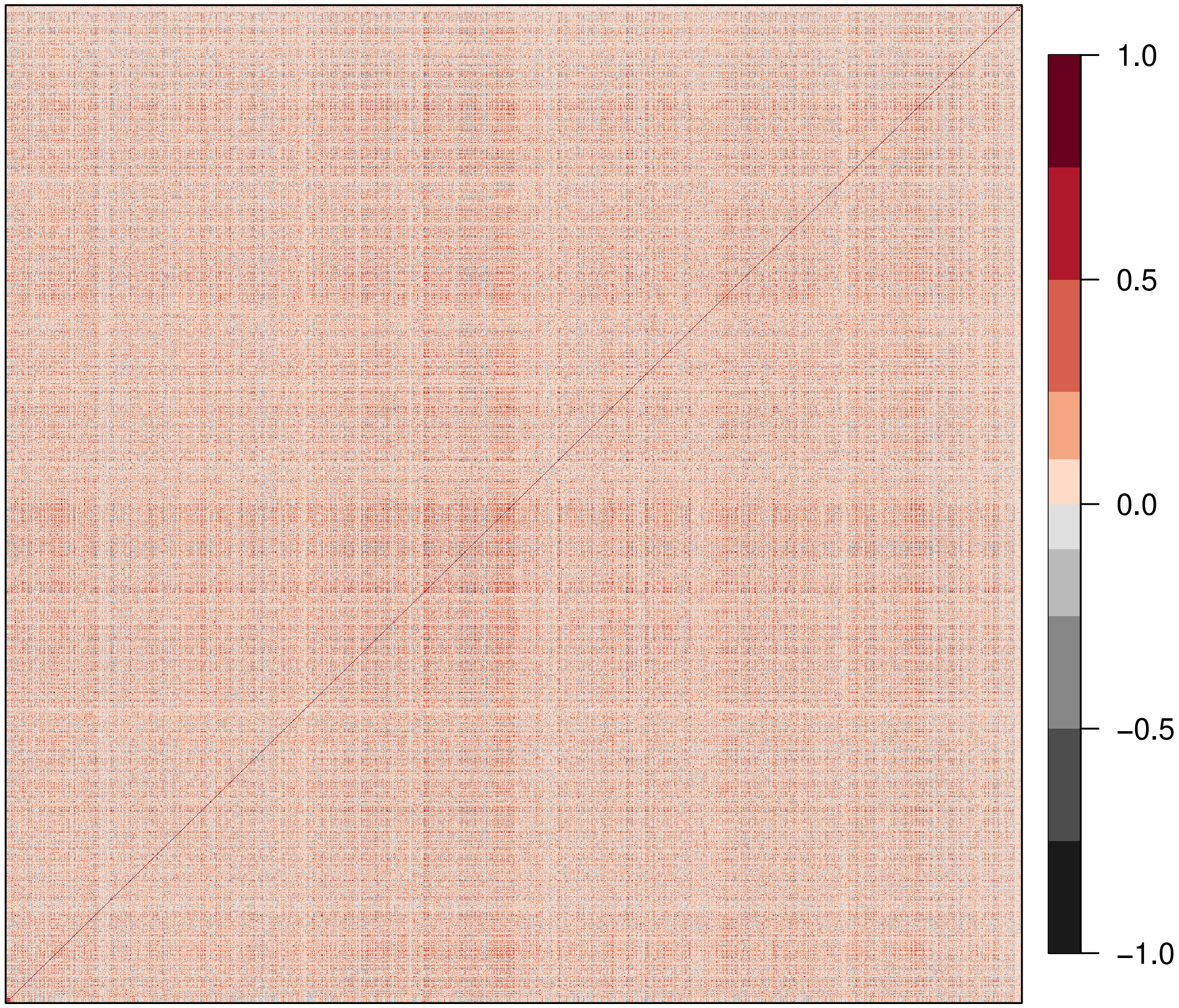


Figure S7: Genetic correlation matrix for the nine caterpillar performance traits (lower left corner), 1750 plant chemistry traits, and 10 other (non-chemical) plant traits (top right corner) based on *M. sativa* genetics from the core common garden. Each colored square in the heatmap denotes the genetic correlation for one pair of traits with the color determined by the Pearson correlation (see the scale on the plot).

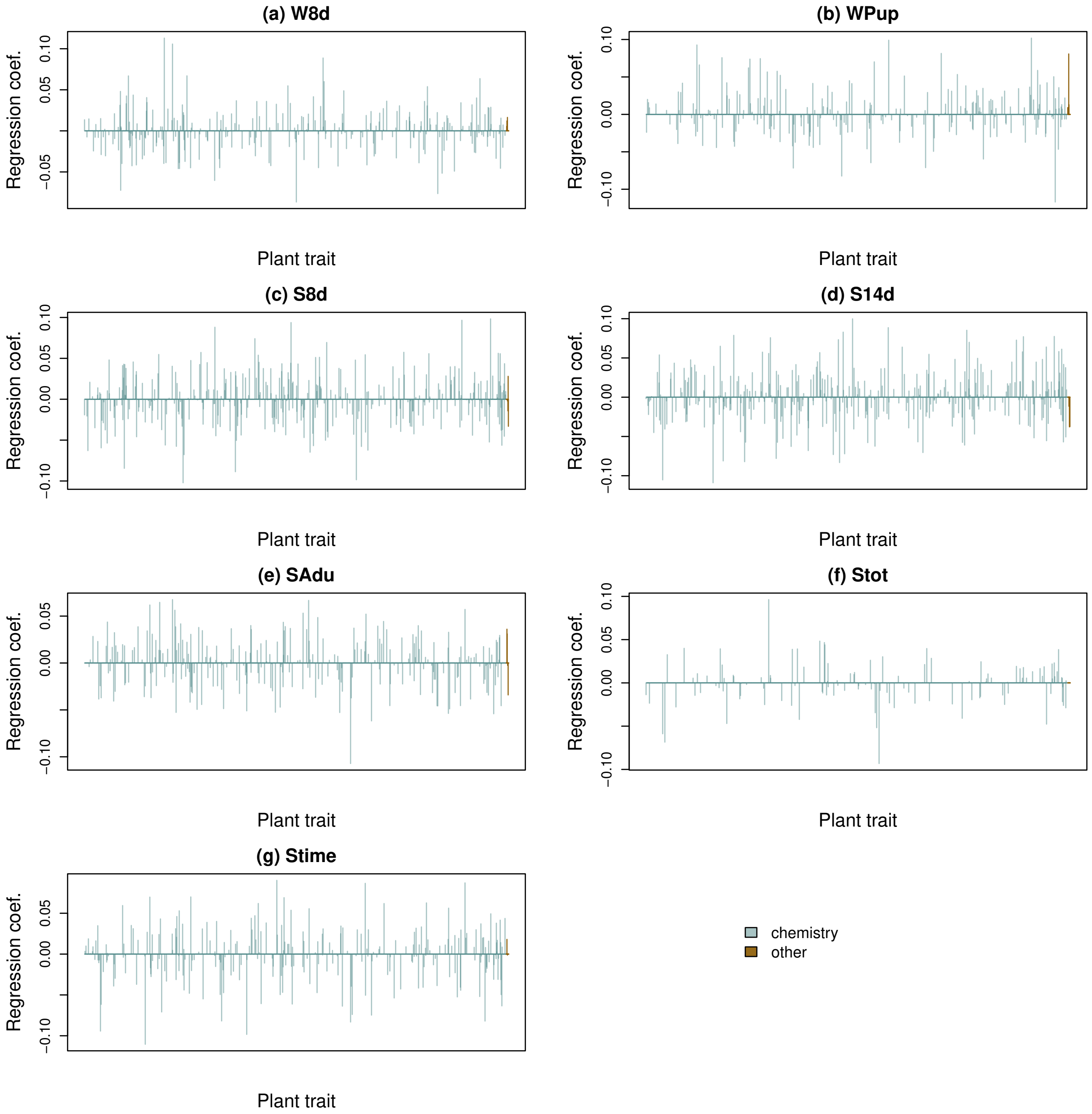


Figure S8: Standardized regression coefficients from LASSO models of caterpillar-performance polygenic scores inferred from *M. sativa* genetics as a function of plant-trait polygenic scores. Results are based on 1760 plant traits and are shown for W8d = 8-day weight, W14d = 14-day weight, Wpup = pupal weight, S8d = 8-day survival, S14d = 14-day survival, SPup = survival to pupation, SAdu = survival to adult, Stot = total survival time, and Stime = (truncated) survival time.

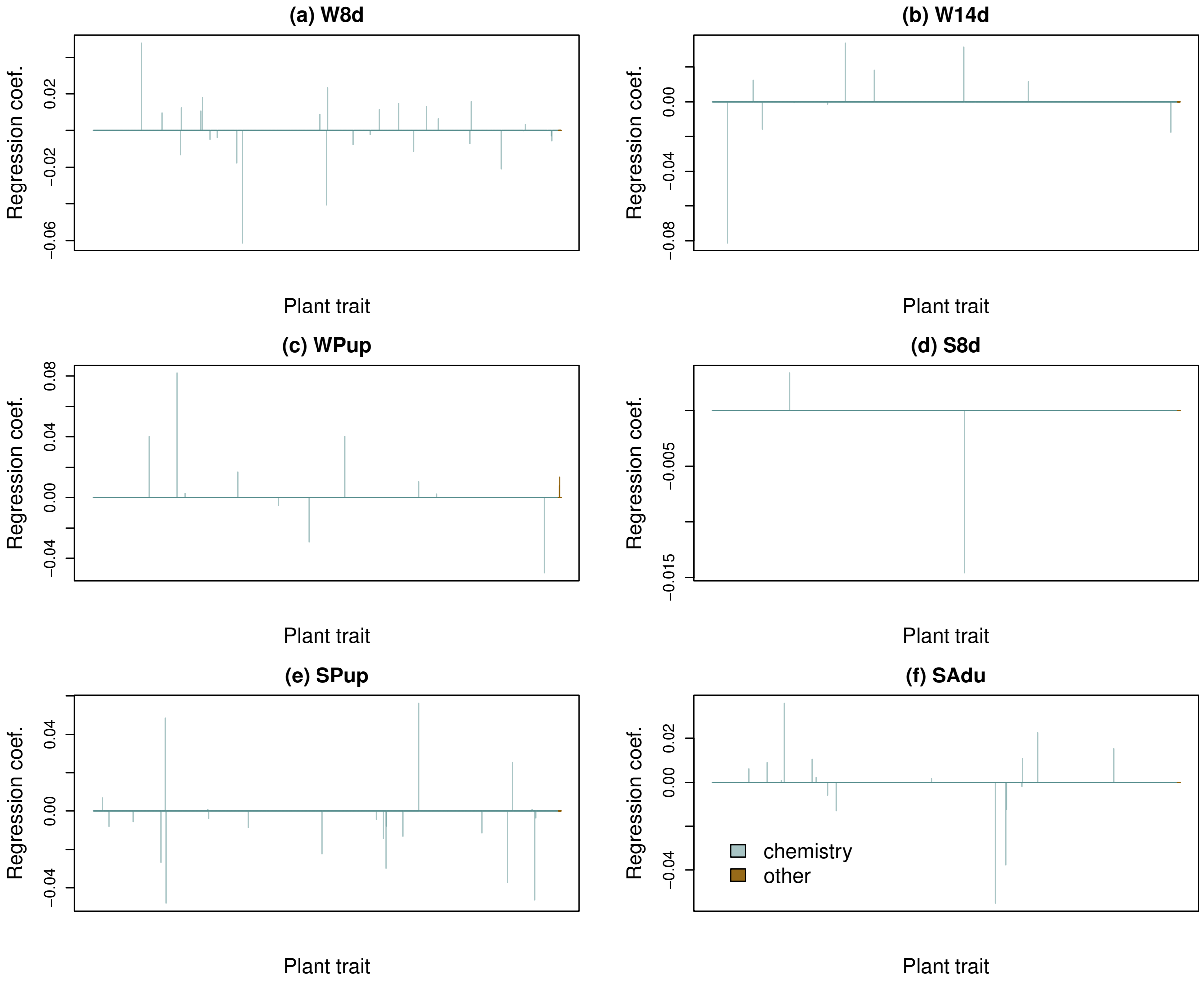


Figure S9: Standardized regression coefficients from LASSO models of caterpillar-performance phenotypes as a function of plant-trait polygenic scores. Results are based on 1760 plant traits and are shown for W8d = 8-day weight, W14d = 14-day weight, Wpup = pupal weight, S8d = 8-day survival, SPup = survival to pupation, and SAdu = survival to adult (no covaraites were retained for the other traits).

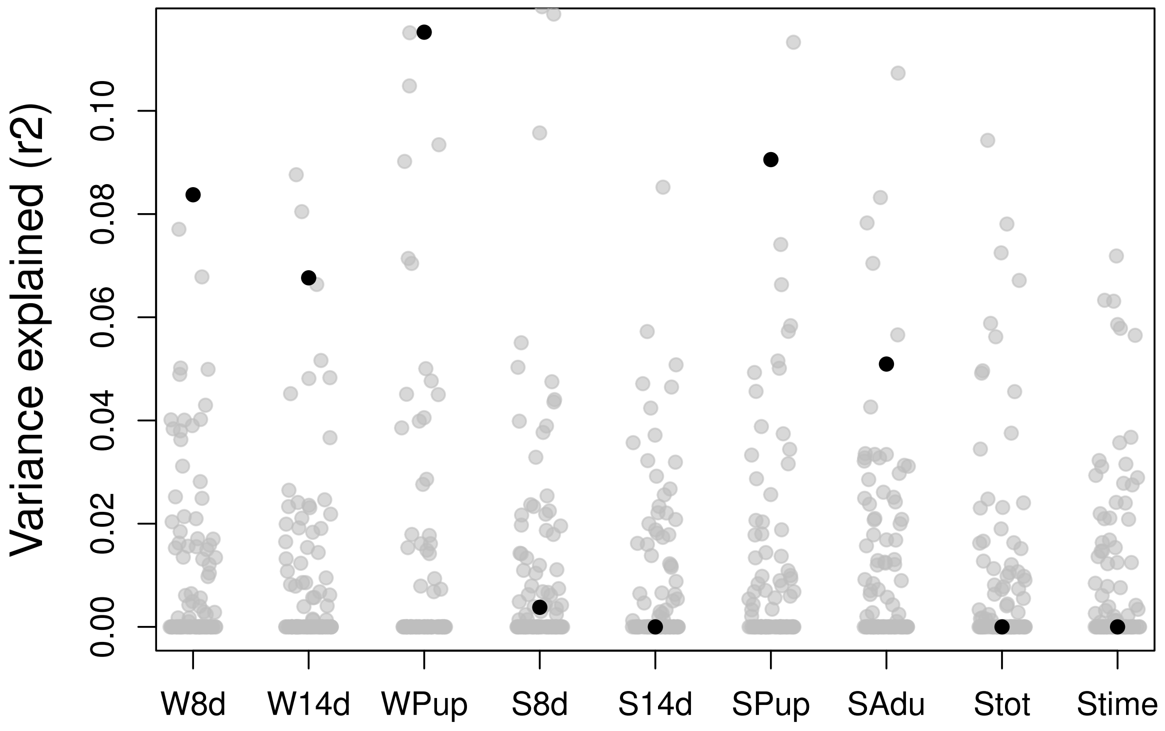


Figure S10: Plot shows the proportion of variance explained in the caterpillar performance traits by LASSO regression models with polygenic scores from 1760 plant traits as possible covariates. Black dotes denote estimates for the observed data and gray dots denote estimates for 100 randomizations of each performance data set. Results are shown for W8d = 8-day weight, W14d = 14-day weight, Wpup = pupal weight, S8d = 8-day survival, S14d = 14-day survival, SPup = survival to pupation, SAdu = survival to adult, Stot = total survival time, and Stime = (truncated) survival time.

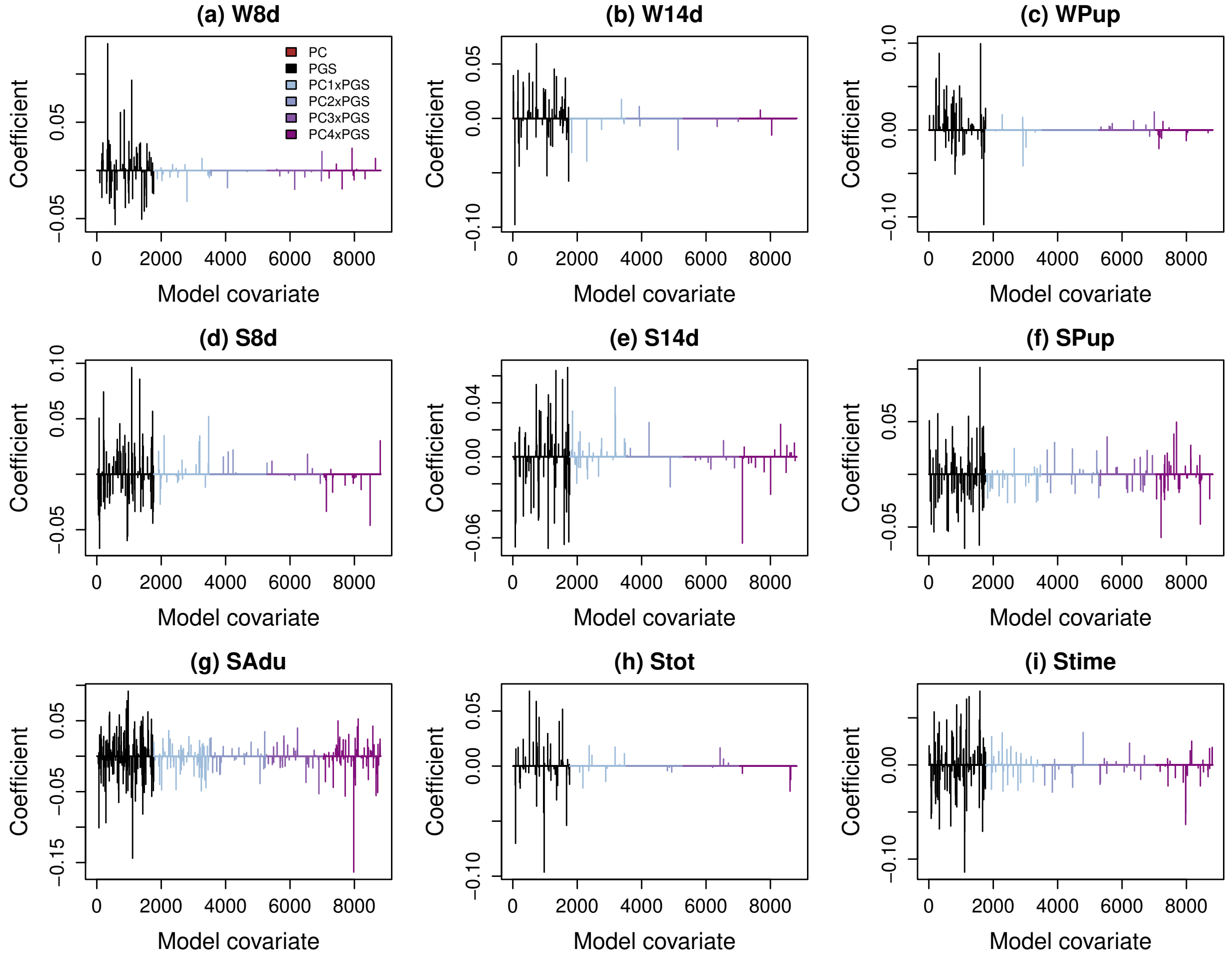


Figure S11: Standardized regression coefficients from LASSO models of caterpillar-performance polygenic scores inferred from *M. sativa* genetics as a function of caterpillar genotype (genetic PCs 1-4) (PC), plant-trait polygenic scores (PGS), and caterpillar genotype-plant trait interactions (PC x PGS). Results are based on 1760 plant traits and are shown for W8d = 8-day weight, W14d = 14-day weight, Wpup = pupal weight, S8d = 8-day survival, S14d = 14-day survival, SPup = survival to pupation, SAdu = survival to adult, Stot = total survival time, and Stime = (truncated) survival time.

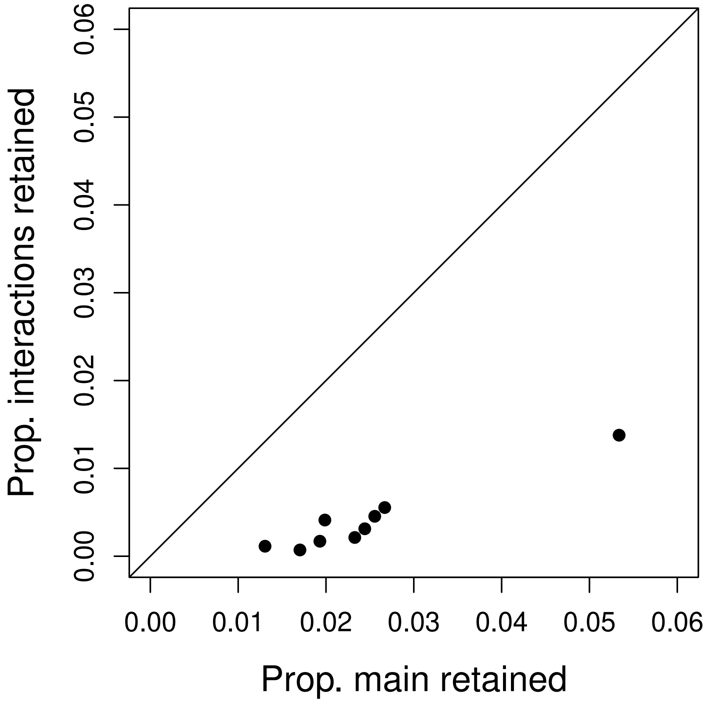


Figure S12: Scatterplot shows the proportion of main effect (non-interaction) versus interaction covariates retained with non-zero regression coefficients from the LASSO regression models of caterpillar-performance polygenic scores as a function of the plant-trait polygenic scores and plant-trait-by-caterpillar genetic PC polygenic scores. A 1:1 line is shown for reference. Each point corresponds with one of the nine caterpillar performance traits.

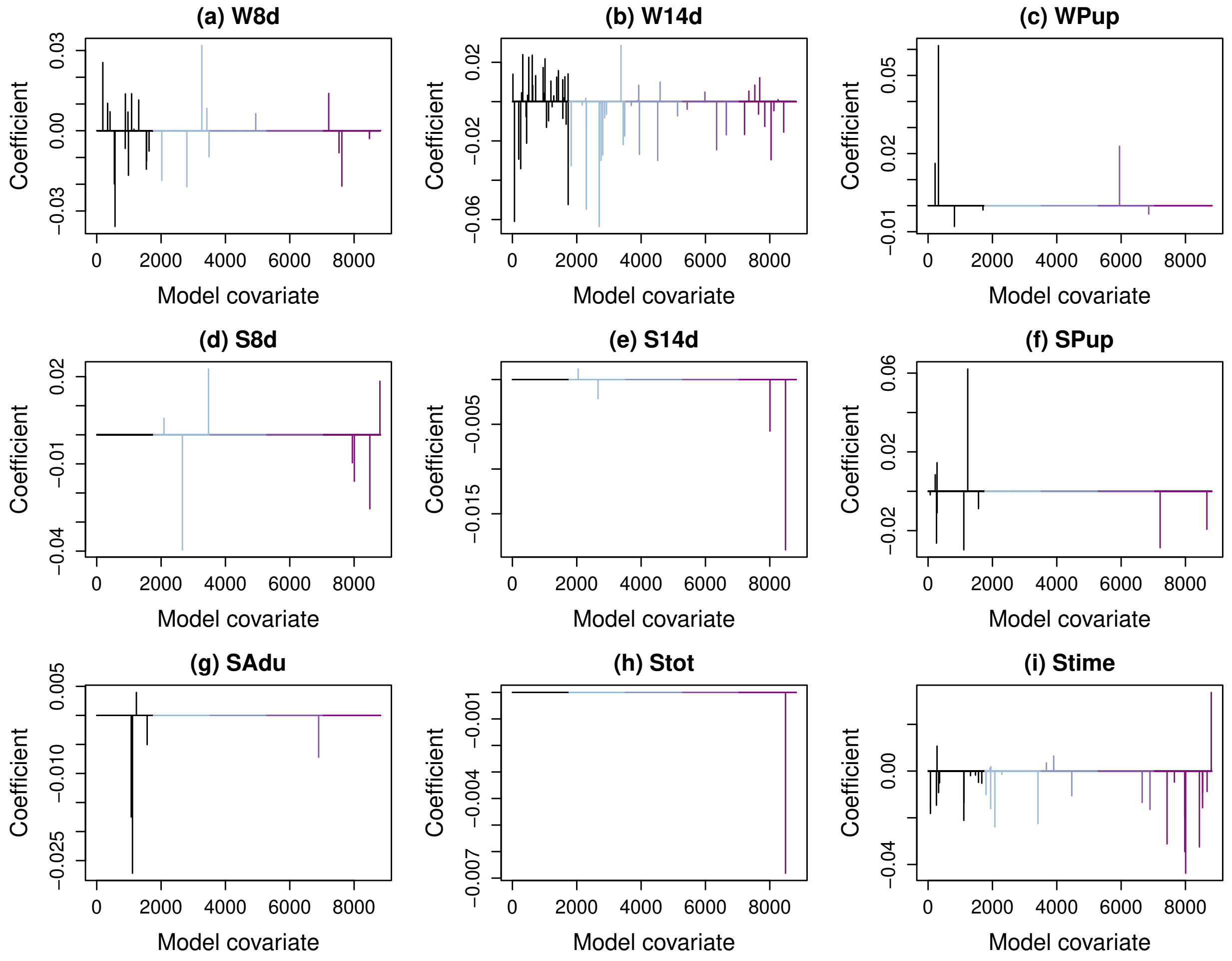


Figure S13: Standardized regression coefficients from LASSO models of caterpillar-performance trait values as a function of caterpillar genotype (genetic PCs 1-4) (PC), plant-trait polygenic scores (PGS), and caterpillar genotype-plant trait interactions (PC x PGS). Results are based on 1760 plant traits and are shown for W8d = 8-day weight, W14d = 14-day weight, Wpup = pupal weight, S8d = 8-day survival, S14d = 14-day survival, SPup = survival to pupation, SAdu = survival to adult, Stot = total survival time, and Stime = (truncated) survival time.

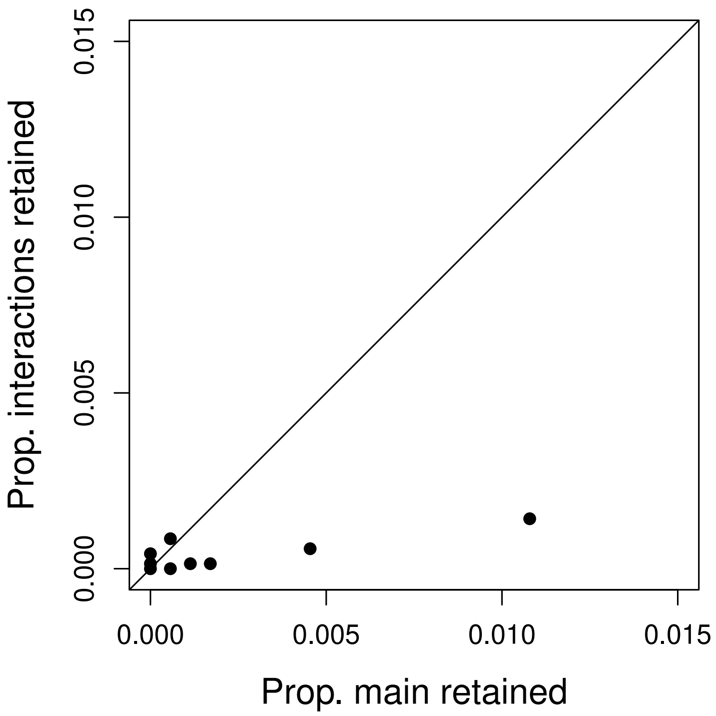


Figure S14: Scatterplot shows the proportion of main effect (non-interaction) versus interaction covariates retained with non-zero regression coefficients from the LASSO regression models of caterpillar-performance trait values as a function of the plant-trait polygenic scores and plant-trait-by-caterpillar genetic PC polygenic scores. A 1:1 line is shown for reference. Each point corresponds with one of the nine caterpillar performance traits.

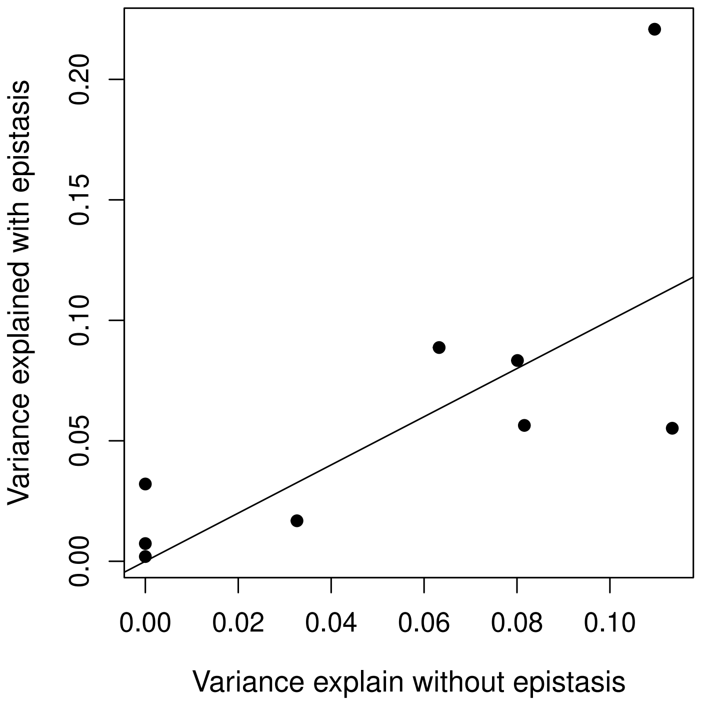


Figure S15: Variance in caterpillar-performance trait values explained by LASSO regression models with versus without epistasis, that is with versus with interactions between plant-trait polygenic scores and caterpillar genetics as captured by genetic PCs. A 1:1 line is shown for reference. Each point corresponds with one of the nine caterpillar performance traits. Although some traits were better explained by the model with epistasis, we found no evidence of an overall effect across traits. Specifically, a linear model of the variance explained with epistasis versus the variance explained without epistasis had an intercept not different from 0 ($\beta = 0.008$, s.e. $= 0.026$, $P = 0.782$) and a slope of $\sim$1 ($\beta = 1.03$, s.e. $= 0.381$, $P = 0.031$, $r^2 = 0.511$).

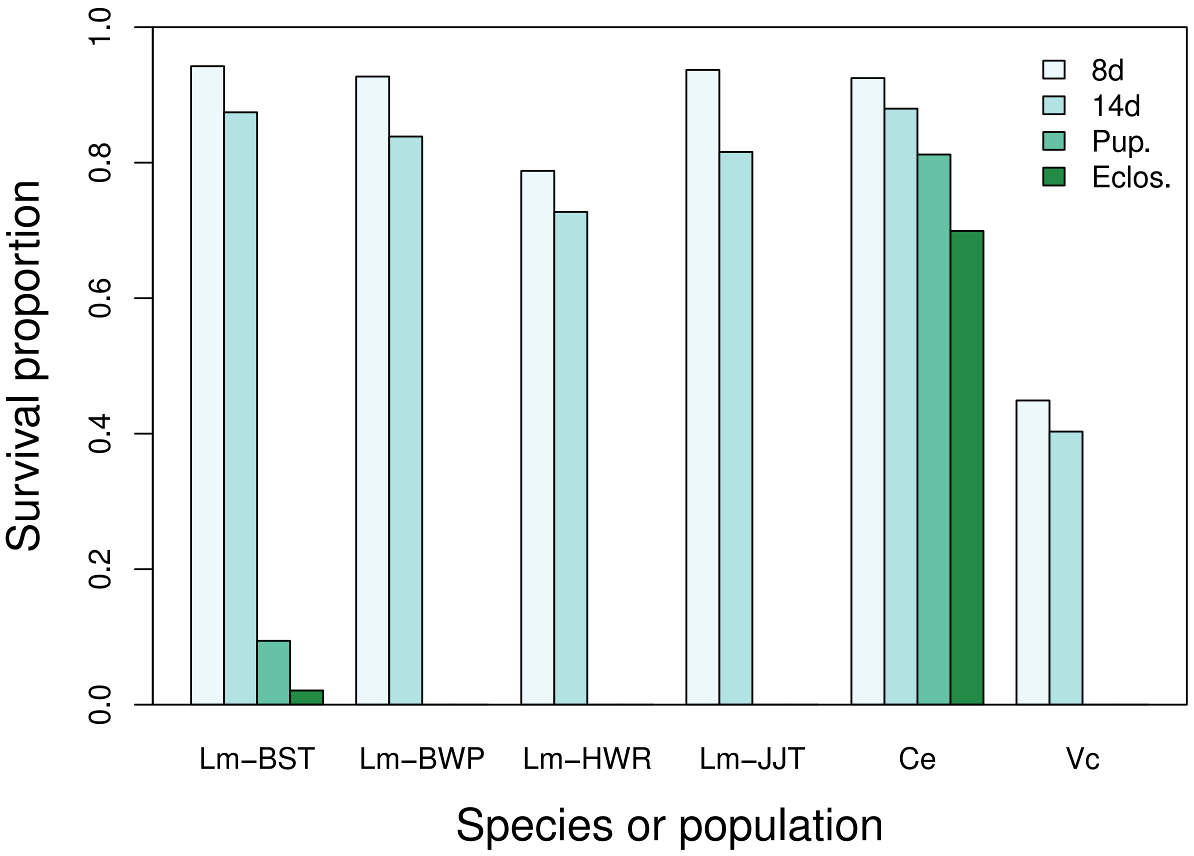


Figure S16: Proportion of caterpillars surviving to 8 days (8d), 14 days (14d), pupation (Pup.), and ecolosion (Eclos.) in the 2018 greenhouse experiment at Utah State University. Results are shown for four *L. melissa* populations (Lm-BST, Lm-BWP, Lm-HWR and Lm-JJT), the orange sulphur (*Colias eurhtyeme*; Ce), which is an alfalfa specialist, and the painted lady (*Vanessa cardui*; Vc), which is a generalist that rarely feeds on alfalfa.

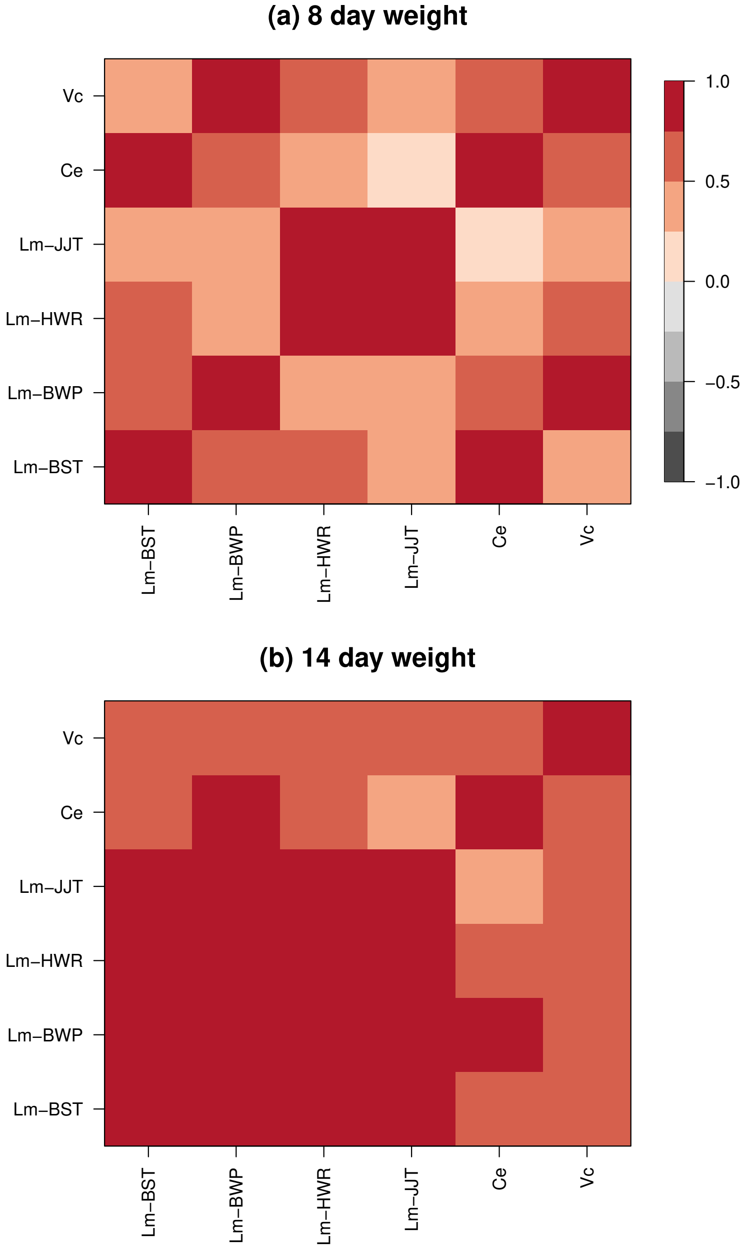


Figure S17: Heatmaps show Pearson correlations across butterfly species and populations for mean 8-day (a) or 14-day (b) weight when caterpillars were fed plants from different greenhouse-grown *M. sativa.* Positive correlations indicate that consistency in plant-population effects on weight across butterfly species or populations. Results are shown for four *L. melissa* populations (Lm-BST, Lm-BWP, Lm-HWR and Lm-JJT), the orange sulphur (*Colias eurhtyeme*; Ce), which is an alfalfa specialist, and the painted lady (*Vanessa cardui*; Vc), which is a generalist that rarely feeds on alfalfa. All 14-day weight correlations between *L. melissa* population pairs were significantly $> 0$ (i.e., $P < 0.05$), along with 8-day weights for *L. melissa* HWR vs JJT, *L. melissa* BST vs *C. eurytheme*, and *L. melissa* BWP vs *V. cardui*; all point estimates of correlations were positive.

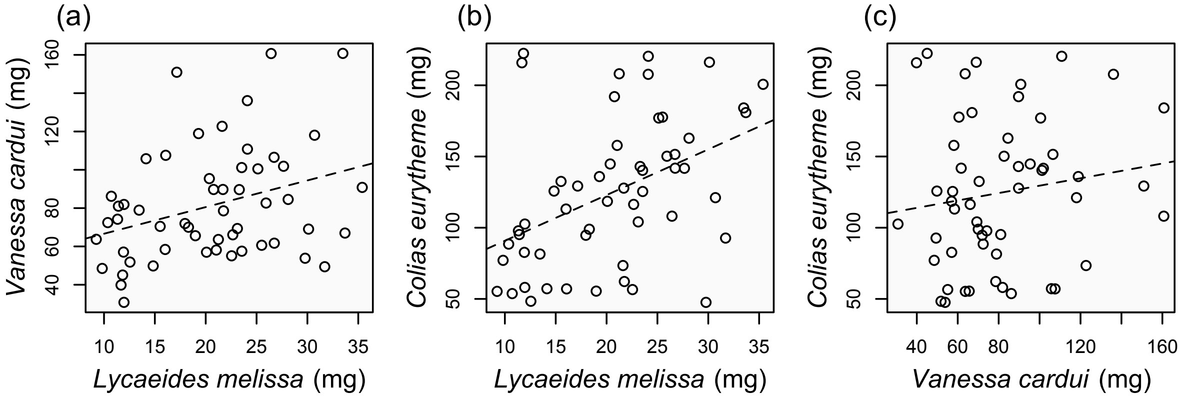


Figure S18: Scatterplots show weight measurements for caterpillars from different butterfly species reared on the same *M. sativa* plant. Best fit lines are shown in each panel.
